# Intrinsically disordered membrane anchors of Rheb, RhoA and DiRas3 small GTPases: Molecular dynamics, membrane organization, and interactions

**DOI:** 10.1101/2024.04.25.591151

**Authors:** Chase M. Hutchins, Alemayehu A. Gorfe

## Abstract

Protein structure has been well established to play a key role in determining function; however, intrinsically disordered proteins and regions (IDPs and IDRs) defy this paradigm. IDPs and IDRs exist as an ensemble of structures rather than a stable 3D structure yet play essential roles in many cell signaling processes. Nearly all Ras Superfamily GTPases are tethered to membranes by a lipid tail at the end of a flexible IDR. The sequence of these IDRs are key determinants of membrane localization, and interactions between the IDR and the membrane have been shown to affect signaling in RAS proteins through modulation of dynamic membrane organization. Here we utilized atomistic molecular dynamics simulations to study the membrane interactions, conformational dynamics, and lipid sorting of three IDRs from small GTPases Rheb, RhoA and DiRas3 in model membranes representing their physiological target membranes. We found that complementarity between lipidated IDR sequence and target membrane lipid composition is a determinant of conformational plasticity. We also show that electrostatic interactions between anionic lipids and basic residues on IDRs generate semi-stable conformational sub-states, and a lack of these residues leads to greater conformational diversity. Finally, we show that small GTPase IDRs with a polybasic domain alter local lipid composition by segregating anionic membrane lipids, and, in some cases, excluding other lipids from their immediate proximity in favor of anionic lipids.

## Introduction

Intrinsically disordered regions (IDRs) of proteins are characterized by functionally relevant rapid transitions among structural ensembles^1,2^. IDRs are most common in proteins involved in signal transduction pathways and membrane binding^3,4^. One example of membrane proteins in which IDRs are indispensable for function is the RAS superfamily small GTPases. RAS GTPases are ubiquitously expressed in mammalian tissues and mediate diverse cell-signaling pathways including the MAPK pathway^5^. Small GTPases are active when GTP-bound and inactive when GTP is hydrolyzed to GDP^6–8^. Dysregulation of this GTPase cycle due to mutation or overexpression leads to many diseases including cancer and developmental disorders^9,10^. The structure of small GTPases is characterized by a highly conserved globular catalytic domain that is tethered to a target membrane by a hypervariable myristoylated N- or prenylated C-terminal IDR^11,12^.

In addition to prenly or myristoyl lipid modification, the IDR of small GTPases often contains a secondary membrane targeting motif such as a polybasic domain (PBD), an amphipathic helix, or an additional lipid modification(s) near the site of penylation or myristoylation. The interplay between these membrane targeting signals and structural plasticity of the IDR determine cellular localization and lipid selectivity^13–15^, as well as membrane orientation dynamics^16–18^. Membrane reorientation is functionally relevant as certain orientations can be deficient in signaling due to occlusion of key effector interacting loop regions by the membrane. Moreover, we have shown previously that the lipid anchor of KRAS, one of the three RAS isoforms in humans, engages in electrostatic interactions with anionic lipids to stabilize specific conformational sub-states^19,20^. These observations highlight how lipidated IDRs play important roles in the signaling function of small GTPases at membrane surfaces.

Despite these progresses, more work is required to elucidate all the shared and unique features of the many diverse lipidated IDRs and how they engage lipids in their respective target membranes. Are there unifying principles underlying the membrane organization of IDR-containing lipidated proteins at membrane surfaces? As part of a broader effort toward addressing this question, we conducted microsecond-scale (∼35 µs aggregate time) atomistic molecular dynamics (MD) simulations of the lipid anchors of three small GTPases from the RAS superfamily: Rheb, RhoA and DiRas3. Since these proteins differ in cellular localization and function, the simulations were conducted in bilayers modeling the presumed target membrane of each protein: RhoA and DiRas3 are primarily localized at the plasma membrane while Rheb is predominantly found at the endoplasmic reticulum^21^ (ER). Rheb regulates cell growth and proliferation by activating mammalian target of rapamycin complex 1 (mTORC1), RhoA regulates cytoskeleton actin organization, and DiRas3 suppresses tumor growth likely by disrupting KRAS signaling on the plasma membrane (PM)^22–24^. The three membrane anchors also differ in lipid-modification: Rheb and RhoA are C-terminally prenylated while DiRas3 is N-terminally myristoylated. The three membrane anchors we have studied thus represent lipidated IDRs of distinct sequence, structure, and target membrane that have potentially unique functional consequences. We show that these IDRs share structural plasticity and preferential interactions with specific lipid species but diverge in their ability to sample a defined set of conformational sub-states that are stabilized by selective interactions with anionic lipids.

## Methods

### Initial peptide structure and system construction

The amino acid sequence, lipid modification, and initial structure of the C-terminally prenylated lipid anchors of Rheb and RhoA as well as the N-terminally myristoylated membrane anchor of DiRas3 (DNTE) are shown in Fig 1. No experimental structure is available for these peptides but they are expected to be largely unstructured except for DNTE, which was found to have ∼14% helical content in solution using circular dichroism (CD) spectroscopy^25^. Therefore, the 11-residue long Rheb and RhoA were modeled as random coils and starting structure for DNTE was prepared as described in SI. These initial structures were attached to a bilayer by manually inserting at least 5 carbon atoms of the myristic or prenyl lipid tail in the hydrophobic core of the membrane, as described previously^20^. The peptide-bilayer system was then solvated with TIP3P waters^26^ and neutralized by adding Na^+^ ions, resulting in the simulation systems listed in Table 1.

**Figure 1:**
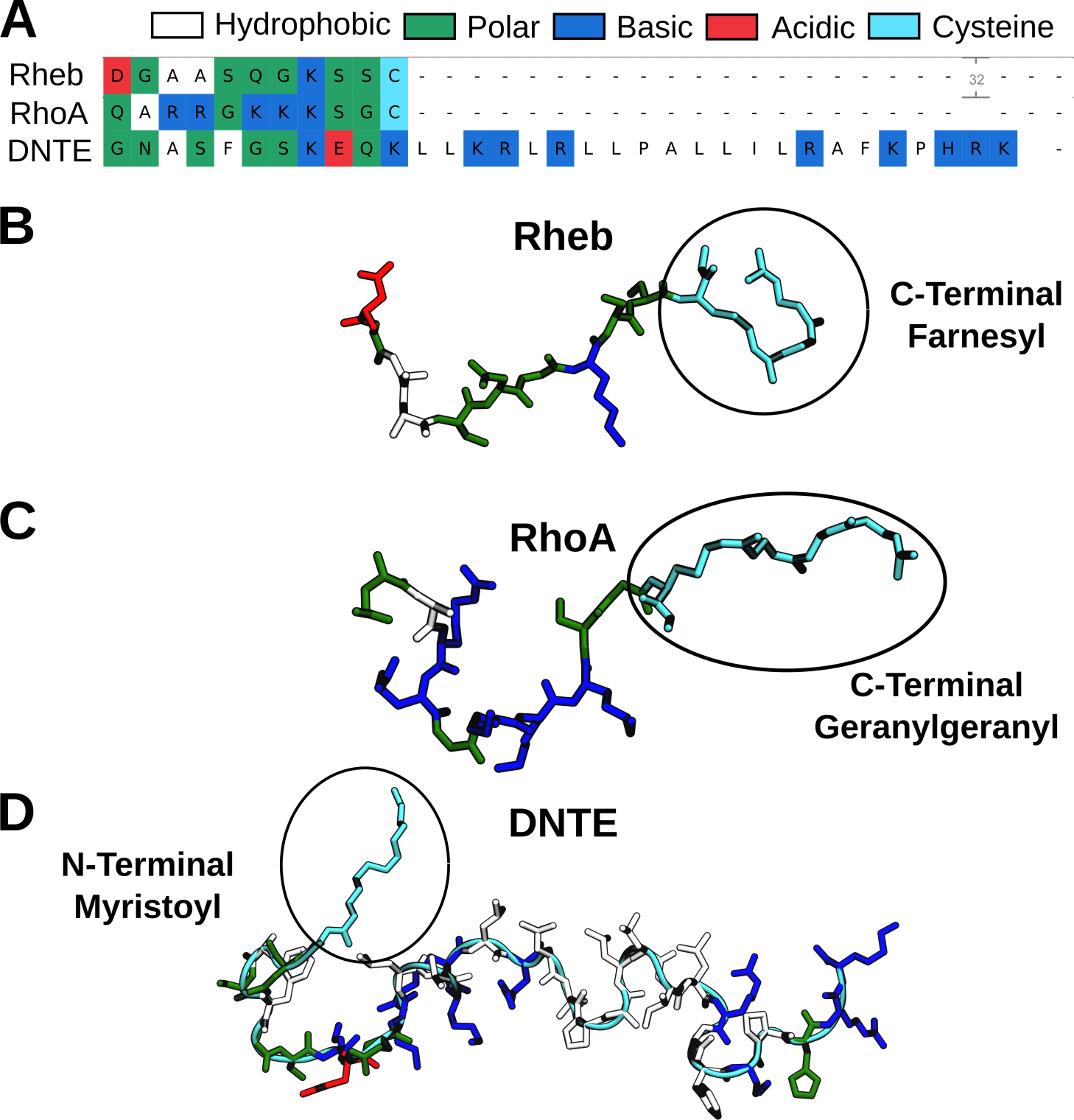
Sequence (**A**) and initial structure (**B-D**) of prenylated C-terminal membrane anchors of Rheb and RhoA and myristoylated N-terminal extension of DiRas3 (DNTE). Color code as indicated.

**Table 1:** Summary of simulations discussed in the main manuscript*.

| Simulation | # atoms | Sim Length (μs) | Lipid Composition | | Bilayer Structure | |
| --- | --- | --- | --- | --- | --- | --- |
| | | | Inner | Outer | Thickness ($\text{\AA}$) | Area ($\text{\AA}^2$) |
| **Rheb** | 57731 | 2 | 7:8:30:55<br>(Chol:PI:PE:PC) | 7:8:30:55<br>(Chol:PI:PE:PC) | $40.4 \pm 0.6$ | $58.6 \pm 0.9$ |
| **RhoA** | 66364 | 2 | 20:30:50<br>(PS:PE:PC) | 100<br>(PC) | $39.0 \pm 0.5$ | $63.8 \pm 1.1$ |
| **DNTE** | 57956 | 10 | 25:25:25:25<br>(PC:PE:PS:PlaPE) | 50:50<br>(PC:PSM) | $39.2 \pm 0.5$ | $61.7 \pm 1.0$ |

\*, See *Supplementary Information* for additional simulations of DNTE. Abbreviations: Chol = cholesterol; PI = phosphatidylinositol; PE = phosphatidylethanolamine; PC = phosphatidylcholine; PS = phosphatidylserine; PlaPE = plasmalogen PE; PSM = sphingomyelin. The molecular structure of each lipid type is found in Fig S1. Membrane thickness is measured as the average z-distance between phosphorous atoms at the two leaflets. Area represents the surface area of the peptide-free monolayer normalized by the number of lipids in that monolayer.

### Model membranes and lipid composition

Initial bilayer models were generated using CHARMM-GUI membrane builder^27,28^ with the composition of lipids determined on the basis of a lipidomic analysis of cell membranes^29^. Rheb is primarily localized in the ER^21^. Therefore, we simulated its lipid anchor in a symmetric bilayer approximating the ER lipid composition, with each leaflet containing 55 POPC, 30 POPE, 7 SAPI and 8 Chol lipids (see Table 1 for abbreviations and Fig S1 for lipid chemical structures). RhoA is primarily localized at the inner leaflet of the plasma membrane (PM) that is enriched in phosphatidylserine (PS) and phosphatidylethanolamine (PE) lipids. Therefore, the RhoA lipid anchor was simulated in a simplified PM model bilayer containing 96 POPC lipids in one leaflet (outer) and 50 POPC, 30 POPE and 20 POPS lipids in the other (lower) leaflet, where the peptide is attached. The difference in the number of lipids at the two leaflets ensures monolayer area symmetry to prevent membrane curvature, as described previously^20^. Even though the ratio of PS to total number of lipids is low in this setup, the 20% PS content in the lower leaflet exposes the RhoA peptide to a similar charge density as in our previous simulations^16,30^, allowing us to make direct comparisons. The longer and more complex DiRas3 N-terminal membrane anchor^25,31^, which to our knowledge has not been simulated before, was initially simulated in a simpler model bilayer of asymmetric PC and PS lipids and then in a more complex asymmetric PM mimic based on a recent paper describing specific leaflet compositions^32^. Here, the peptide-free (upper) leaflet contained POPC and PSM lipids of equal ratio while the peptide-containing lower leaflet contained equal numbers of POPC, POPE, PlaPE, and POPS lipids (Table 1).

### MD simulations

All simulations were conducted using the CHARMM36 forcefield^33^. Rheb and RhoA were equilibrated with the NAMD simulation package^34^ and DiRas3 with GROMACS^35^. Similar equilibration protocols were used for all systems, with each system energy-minimized for 2000 steps with lipid phosphorus and protein backbone atoms fixed, and then for another 2000 steps without fixed atoms using the conjugate gradient (NAMD) or the steepest descent (GROMACS) method. After minimization, systems were equilibrated with Δt = 1 fs for 4 ns applying a harmonic restraint of force constant k = 4 kcal/mol Å^2^ to lipid phosphorus and protein backbone atoms. The restraint was scaled down by 25% in four steps of 1 ns each until k = 0. The systems were then simulated for at least 40 ns on local resources with Δt = 2 fs, using the particle mesh Ewald (PME) method^36^ for long-range electrostatic interactions and restraining bonds involving hydrogen atoms with SHAKE^37^ (NAMD) or LINCS^38^ (GROMACS). A switching function with 10 and 12 Å distance cutoffs was used, with a 14 Å cutoff for pair-list update. The NPT (constant number of particles, pressure, and temperature) ensemble was used with the Nose-Hoover Langevin piston and Langevin thermostat (NAMD) or the velocity-rescaling thermostat^39^ and Parinello-Rahman barostat^40^ (GROMACS) to maintain pressure at 1 bar and temperature at 310 K, respectively. After equilibration, RhoA and Rheb were simulated for 40 ns and DNTE for 400 ns on local resources and then transferred to Anton 2^41^. Anton simulations used default Desmond parameters for the NPT ensemble and a 2.5 fs time step and ran for 2 µs (RhoA and Rheb) or 10 µs (DNTE) with snapshots written out every 100 ps for analysis.

### Trajectory analysis

Trajectories were analyzed using the MDAnalysis python library^42^ or GROMACS analysis tools, with additional analysis performed with Visual Molecular Dynamics (VMD) scripts^43^. Statistical analyses were performed in Python using the Scipy library^44^. For Rheb and RhoA, the best equilibrated last 1.5 µs of the trajectories was used to obtain equilibrium properties. We used the entire 10 µs trajectory of DNTE for analysis because this simulation was started from an already well-equilibrated bilayer system (see SI text and Fig S2). Bilayer thickness (P-P) was measured using the z-component of the distance between the centers-of-mass (COM) of the phosphorus atoms in the two leaflets. Area per lipid (APL) was calculated using the area-per-lipid class in the Lipyphilic Python toolkit^45^, which utilizes Voronoi Tesselations to calculate the surface area of individual lipid molecules. Probability density functions were calculated using the Gaussian KDE method from scipy stats. 2D particle densities calculations were performed by wrapping lipid coordinate positions around the COM of each protein and calculating the Gaussian kernel density distribution of X/Y coordinates of lipid phosphorus atoms around the protein COM. We used mean square displacement (MSD) calculations to obtain lateral diffusion coefficient (D) of lipids. We have used multiple reaction coordinates for structural analyses of the peptides. These included end-to-end distance (*d*) of the backbone to measure compactness, pseudo dihedral angles (*a*) defined by four consecutive Cα atoms to assess local structural features, and pair-wise distances between non-adjacent Cα atoms to perform principal component analysis (PCA) with the PyEMMA library^46^. Secondary structure content was calculated using the DSSP^47^ method implemented in the MDTraj Python library^48^. Peptide-lipid interactions were monitored by counting the number of hydrogen bonds (defined by an angle cutoff of 130° and distance cutoff of 3 Å) and van der Waals (vdW) contacts (defined to occur if two carbon atoms are within 4 Å of each other).

## Results and Discussion

The current work has focused on MD simulation analyses of the lipidated IDRs shown in Fig 1 bound to the model membranes listed in Table 1. The time evolution of bilayer thicknesses (Fig S3) and surface areas (not shown) indicated that all three bilayers have fully stabilized during the equilibration phase of the simulations, fluctuating around the mean values listed in Table 1. Plots of backbone root-mean-square deviations (RMSD) and membrane insertion depth of the lipid-modified residue (I_D_), as well as the numbers of hydrogen bonding (N_HB_) and vdW (N_vdW_) contacts (Fig S4) show that all three peptides quickly adsorbed and stabilized in their respective bilayers (e.g., within 0.5 µs in RhoA and Rheb). Based on the RMSD profiles, we decided to use the last 1.5 µs data of Rheb and RhoA and the entire trajectory of the more complex DNTE for analysis of equilibrium properties.

A bilayer similar to our RhoA model membrane has been simulated before^20^ but, to our knowledge, the Rheb and DNTE bilayer systems were not previously studied with simulations. Therefore, we first briefly discuss the structure and dynamics of the three bilayers based on typical structural properties such as bilayer thickness (P-P distance), APL, diffusion coefficient (D), and acyl chain order parameters. We will then describe our findings on the membrane interaction, structure, and dynamics of the lipidated IDRs.

### Bilayer structure and dynamics

Table 1 shows that the P-P distance in the RhoA and DNTE bilayers is 39.0 and 39.2 Å, respectively. Their peptide-free monolayer surface area normalized by number of lipids in that monolayer is also comparable (Table 1). These suggest that, despite differences in lipid composition, our PM models are structurally similar to each other and to other PM model membranes such as PC-PC/PS and PC-PC/PS/PE bilayers^20,49^. By contrast, an average thickness of 40.4 Å in the Rheb bilayer indicates that our ER model membrane is slightly thicker than the PM models. This is expected because cholesterol increases lipid packing and bilayer thickness and decreases area per lipid^50,51^. Consistent with the trend in bilayer thickness (Table 1), the APL of both POPC ((APL(PC)) and POPE (APL(PE)) is slightly smaller in Rheb than in RhoA or DNTE (Fig 2A). This is consistent with C^13^-based nuclear magnetic resonance (NMR) experiments showing that cholesterol reduces APL of a POPC bilayer^50^. APLs of SAPI and Chol from the Rheb simulation (Fig 2A) are consistent with expectation from the size of their head group (Fig S1).

**Figure 2:**
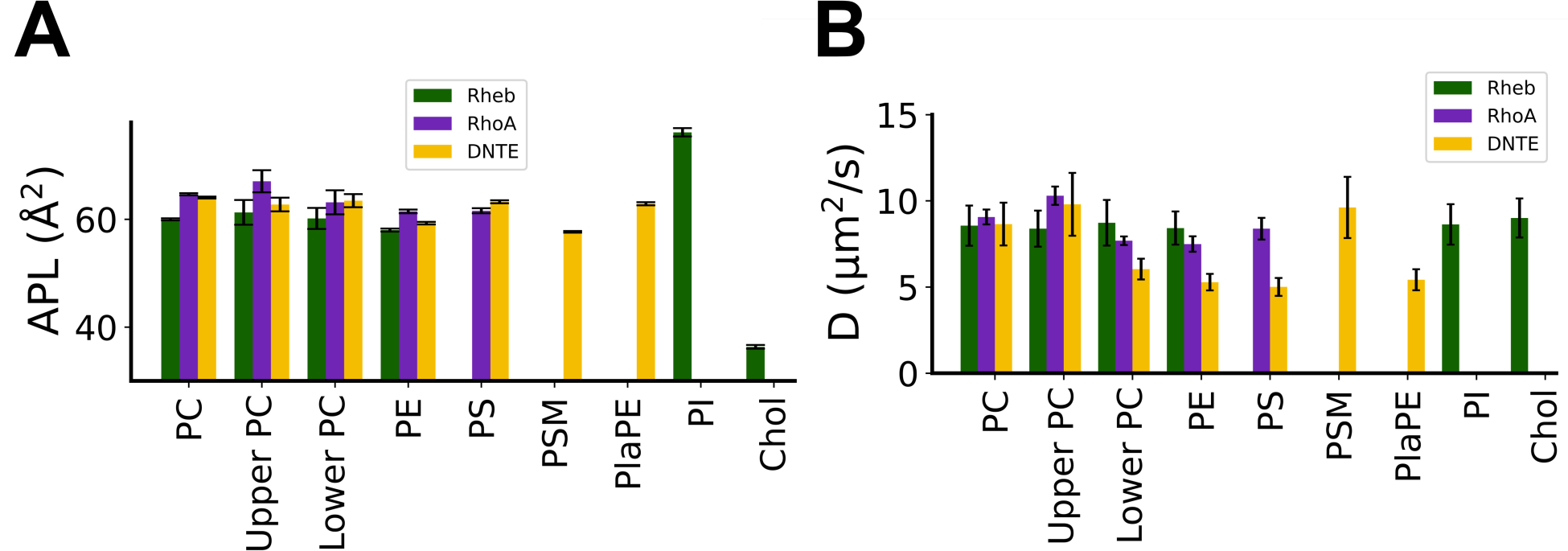
Time averaged area per lipid (APL; **A**) and lateral diffusion coefficient (D; **B**) of each lipid type in each simulation. APL was calculated using the last 1.5μs of Rheb and RhoA simulations and the entire trajectory of DNTE. D was obtained from mean square displacements (MSD) calculated using 200 ns time blocks of the last 1 μs data, with linear regression performed on the most linear portion of the MSD plots, as described previously^20^.

We obtained APL = 67.1 ± 2.0 Å^2^ for POPC in the peptide-free leaflet of the RhoA simulation, a value similar to that from a previous simulation of a similar mixed bilayer (66.2 ± 1.3 Å^2^) ^20^, and experimental data in a pure POPC bilayer (67.3 Å^2^ at 323 K)^52^. APL(PC) in the peptide-containing lower leaflet of RhoA is slightly smaller than for the upper leaflet likely as a consequence of interactions with the bound peptide. APL(PS) and APL(PE) from the RhoA simulation are comparable to data from a previous simulation of the same lipid mixture^20^. As in the P-P distances, the average APL of lipids in the compositionally more complex DNTE bilayer is similar to those in the simpler PM mimic RhoA. However, there is no difference in APL(PC) between leaflets, and APL(PS) is slightly larger and APL(PE) slightly smaller in DNTE than in RhoA (Fig 2A). APL(PSM) and APL(PlaEP) are consistent with expectations from molecular structures (Fig S1).

Acyl chain order parameters (S_CH_) plotted in Fig S5 show that the three bilayers are remarkably similar in lipid packing despite differences in composition and symmetry. Still, both the Sn-1 and Sn-2 acyl chains of PC and PS lipids are somewhat more ordered in Rheb than in the other bilayers (Fig S5), consistent with the increased bilayer thickness and reduced APL discussed above. The S_CH_ profiles of each lipid type in the simpler RhoA and more complex DNTE PM mimics are very close to each other. Moreover, acyl chain dynamics are identical between the peptide-containing lower and peptide-free upper leaflets in RhoA and DNTE. The S_CH_ profiles of lipids found only in Rheb (PI) or DNTE (PSM and PlaPE) do not display any unexpected behavior. In the RhoA bilayer, POPC acyl chains in the peptide-containing lower leaflet are slightly less ordered than in the upper leaflet, reflecting the inter-leaflet difference in APL(PC) and reproducing a previous observation in a similar bilayer with KRAS membrane anchor^20^. Overall, the two compositionally different PM mimics are characterized by very similar average structural properties that differ from our ER model membrane Rheb.

To assess the consequence of these structural differences and similarities in lipid dynamics, we calculated the lateral diffusion coefficient, D, of each lipid type (Fig 2B). It is immediately clear that lipids in the DNTE membrane are generally less mobile than in Rheb and RhoA, and that the latter two share similarity in terms of lipid mobility (Fig 2B). For POPC, we obtained an average D ≈ 9.0 µm^2^/s in all simulations, a value close to previous observations from simulations^20,53^ and experiments^54^. Similar values were obtained for POPS, POPE, SAPI, and Chol in the RhoA and Rheb simulations and for PSM in DNTE (Fig 2B). However, we obtained a lower value of D ≈ 6.0 µm^2^/s for POPS, POPE and PlaPE lipids in the DNTE system. Lipids including POPC in the peptide-containing lower leaflet are less mobile in RhoA and even more significantly in DNTE. The restricted lateral dynamics of lipids in the lower leaflet is likely due to interaction with the bound peptide, which will be discussed in subsequent sections.

### Rheb membrane anchor is flexible and RhoA samples multiple semi-stable conformational sub-states

As noted above and shown in Fig S4B using the membrane insertion depth of the prenyl chains (I_D_), both Rheb and RhoA rapidly inserted and stabilized in their respective bilayer. Following insertion, RhoA fully adsorbed into the bilayer with the backbone lying flat in the head group region (Fig 3). In contrast, Rheb remained extended and flexible, interacting with the host monolayer primarily via its farnesyl chain (Fig 3). These observations are consistent with previous findings from simulations in asymmetric PC-PC/PS bilayers^49^, with the difference between the two peptides in bilayer localization ascribed to RhoA’s ability to engage anionic lipids through its PBD. This is supported by the 2-fold higher number of peptide-lipid HB interactions (N_HB_) in RhoA (5.5 ± 2.4) than in Rheb (2.6 ± 1.5) (Fig 4A & S4D). Unsurprisingly given the PBD, RhoA predominantly interacts with PS (and PC to some extent) while Rheb interchangeably and weakly engages PC and PE (Fig 4A). The radial pair distribution plots in Fig 4B further show that each of the three Lys and two Arg residues in the RhoA PBD strongly interact with PS and to a lesser extent with PE, whereas the single Lys in Rheb only modestly interacts with POPE. The Rheb and RhoA IDRs also differ in the number of vdW contacts (N_vdW_) they make with lipid acyl chains (Fig S4C). We obtained N_vdW_ = 24.5 ± 9.4 for RhoA and 17 ± 6.9 for Rheb. This difference is almost completely a result of the longer 20-carbon geranyl geranyl acyl chain in RhoA versus the 15-carbon farnesyl in Rheb. Taken together, these results suggest that not only do Rheb and RhoA IDRs differ in structure and membrane adsorption but also in apparent affinity for their host membrane, with RhoA likely attaching to the PM more tightly than Rheb to the ER membrane.

**Figure 3:**
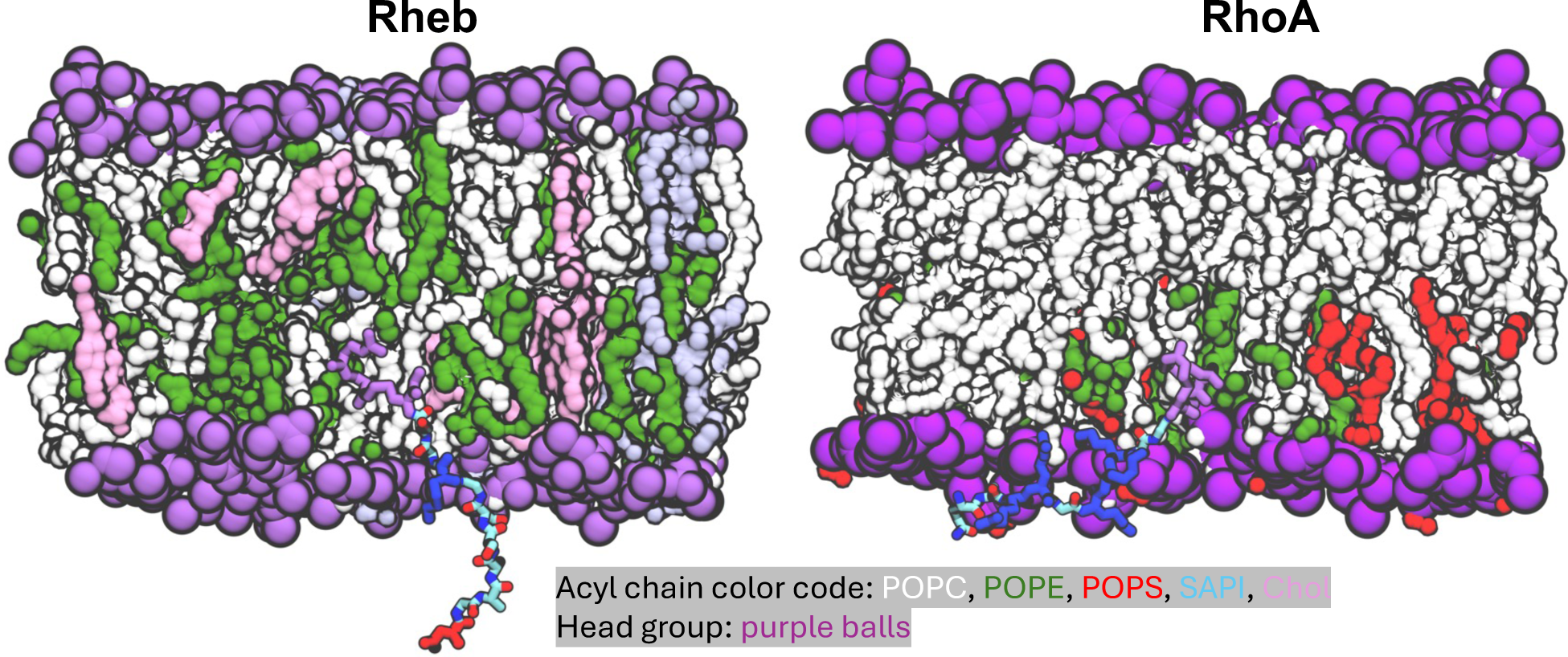
Snapshot at the end of Rheb and RhoA simulations illustrating the organization of the lipidated IDR in bilayer, highlighting basic residues in blue, acidic in red, lipidated cysteines in magenta and remaining backbone atoms in atom-colored licorice. Protein backbone is shown in cartoon and side chains in licorice. Lipid acyl chains are shown as surface and head groups as sphere colored as indicated. Lipid atoms including those of POPS within 10 Å of protein heavy atoms as well as water and ions are omitted for a better visualization of the peptide.

**Figure 4:**
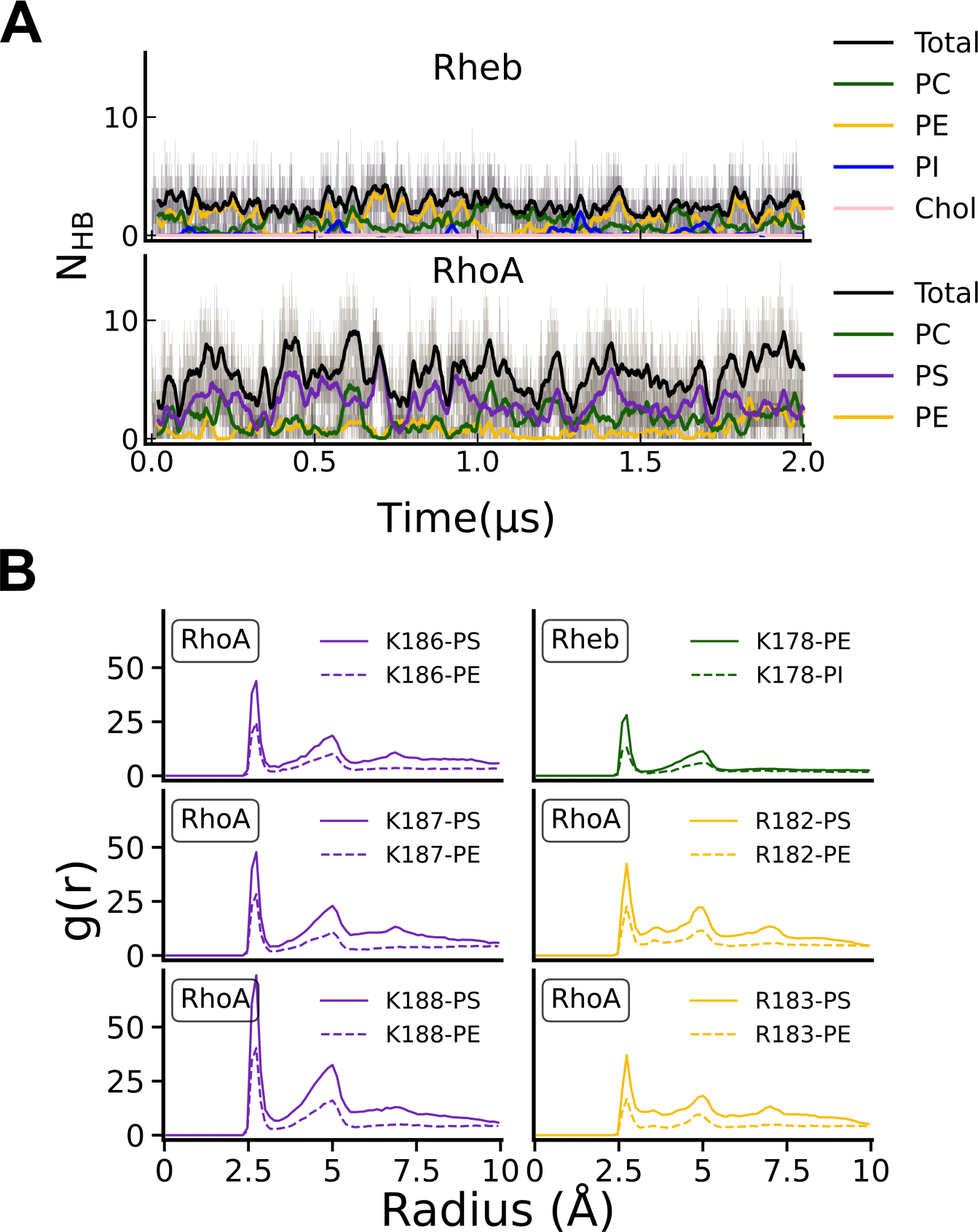
(**A**) Time evolution of the number of hydrogen bonds (N_HB_) between all protein hydrogen bond donors and phosphate oxygen atoms of each lipid type in Rheb (top) and RhoA (bottom) simulations. (**B**) Radial pair distribution of phosphate oxygen atoms around the NZ atom of lysine and NH1 atom of Arg side chains on the Rheb and RhoA anchors.

We have shown in previous studies^19,20,49^ that one consequence of differential bilayer organization and lipid interaction of lipidated IDRs is an altered backbone conformational dynamics. To test if this is the case in the current Rheb and RhoA simulations, we analyzed simulated conformers of each trajectory (last 1.5 µs) using backbone end-to-end distance (*d*) and pseudo-dihedral angle (*a*) as reaction coordinates to respectively measure global and local structural features (see Methods). Plots of *P(d,a)* probability density distributions (Fig 5A) show that Rheb predominantly samples extended conformations of specific planarity, with occasional excursions into conformations of local structure characterized by *a* > 0.0 °. In contrast, RhoA sampled multiple meta-stable states including a compact semi-ordered conformation and extended structures with either *a* < 0.0 ° or *a* > 0.0 °. As discussed later, these meta-stable conformations are stabilized by preferential interactions with anionic lipids (Fig 5B), indicating that interplay between IDR sequence and lipid composition underlies the conformational energy landscape of lipidated IDRs.

**Figure 5:**
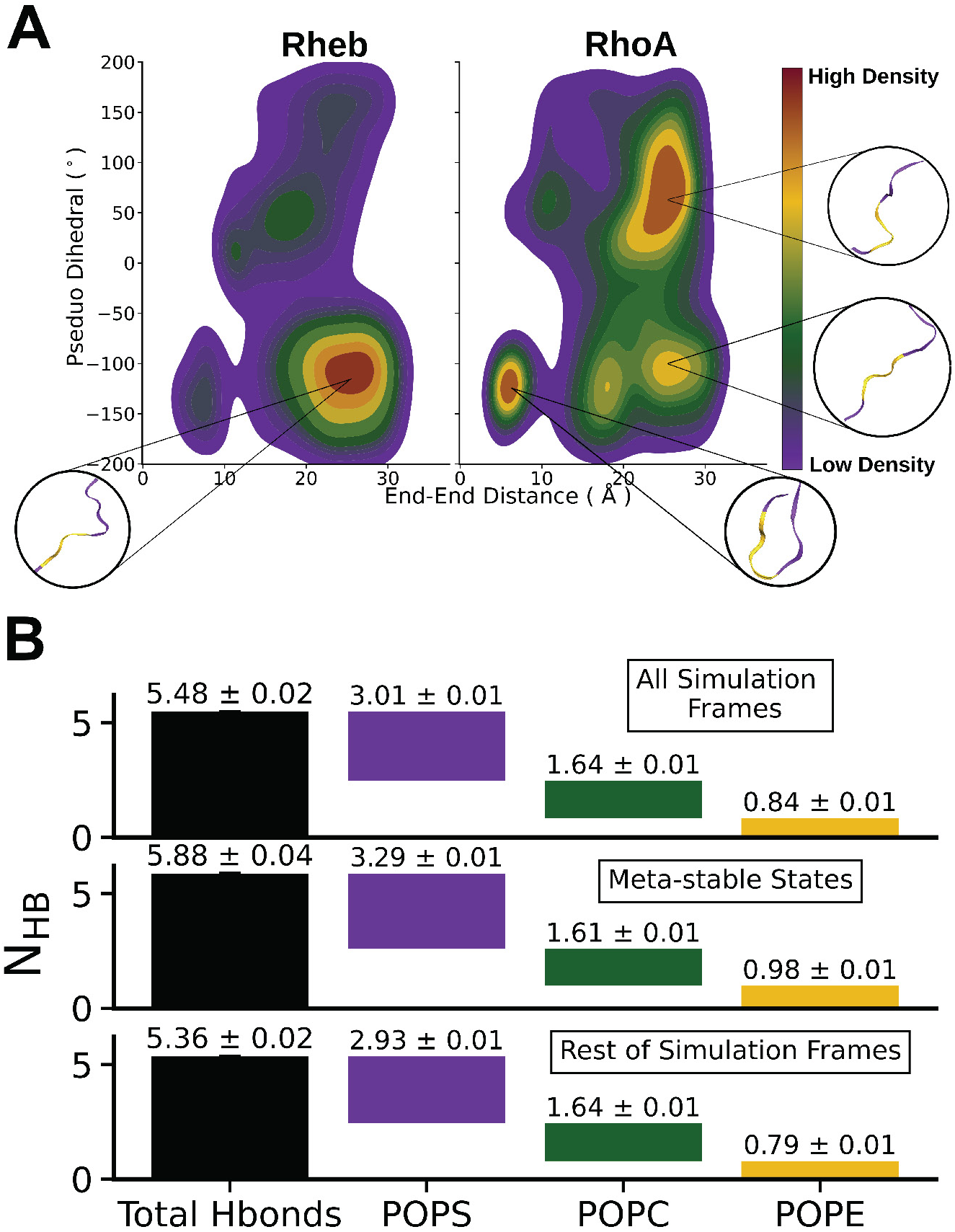
(**A**) Peak-normalized 2D probability number density distribution of the Rheb and RhoA simulated conformers along a pseudo dihedral angle and end-end distance reaction coordinates. The dihedral angles were calculated using four consecutive Cα atoms and the distance was between Cα atoms of C and N-terminal residues. Representative structures for the highest peaks are shown in purple with residues used for pseudo-dihedral angle calculation marked in gold. (**B**) Waterfall plots of average number of hydrogen bonds between RhoA and lipids comparing all semi-stable conformations (combining conformers in all the high-density contours in Fig 2) and the rest of the conformers.

### DNTE undergoes disorder to order transition upon binding to and stabilizing in an anionic membrane

As described in SI, the DNTE simulation discussed here was preceded by extensive (21 µs aggregate time) exploratory simulations in search of an initial structure and setup that may produce reliable results and conforms to known patterns of amphipathic helix membrane binding^55–57^. In this final simulation, we found that DNTE quickly adsorbed and stabilized in the bilayer as in the Rheb and RhoA simulations, with its I_D_ fluctuating around a mean value of ∼11 Å (Fig S4B). However, unlike the smaller Rheb and RhoA peptides, DNTE’s electrostatic and vdW interactions with lipids continued to evolve until about 3 µs, as can be seen from the plateauing of the time evolution of N_HB_ and N_vdW_ after this time point (Fig S4C,D). The backbone RMSD fluctuated until about 5 µs (Fig S4A). Note that the initial structure of DNTE was characterized by a helix-turn-helix motif with the N-terminal **α**-helix (residues 8-16) persisting throughout our exploratory simulations (Fig S2). Therefore, we monitored changes in secondary structure content during the final simulation and found that DNTE undergoes substantial structural changes (Fig S6). This included the N-terminal helix unfolding quickly, including during a 400 ns equilibration run, and the C-terminal helix re-emerging within the first 100-200 ns of production run (Fig S6). The C-terminal helix was unstable until about 3 µs but fully stabilized as a 9 amino acids-long helix (residues 21-29) in the second half of the trajectory, after the stabilization of N_HB_, N_vdW_ and RMSD (Figs S4 & S6). In addition, pi- and 3/10-helices sporadically formed throughout the simulation, and a short helix spanning residues 2-6 appeared at the end of the simulation (Fig S6). In short, DNTE’s helical content fluctuated between 0 and 20% in the first half and stabilized at ∼30% during most of the second half of the trajectory, reaching ∼40% at the end of the simulation. Consistent with this observation, it was found experimentally that DNTE’s helicity increases upon binding to anionic lipids^25^. Moreover, mutating the myristoylatable Gly 2 (first residue in Fig 1) to Ala did not affect helical content or response to lipid binding, and a peptide made up of residues 2-11 of DNTE has a lower helicity than the wild type (8 % versus 14 %) in solution. Importantly, the helical content of this peptide did not increase in the presence of anionic lipids. In contrast, a peptide made up of residues 12-29 of DNTE transitioned from 11% helicity to mostly helical upon lipid binding^25^. That DNTE becomes more helical and its C-terminal, but not N-terminal, portion undergoes disorder to order transition upon binding to anionic lipids validate our simulation results.

### DNTE interacts with lipids extensively through both hydrophobic and basic residues

Like the PBD-containing RhoA membrane anchor, the entire structure of DNTE with the exception of the polar residues around Gln 9 submerged in the bilayer (Fig 6A,B). Note that while polar or basic amino acids dominate in Rheb and RhoA membrane anchors, DNTE harbors mostly polar (residues 1-10, which we call p) and hydrophobic (residues 18-25, np) segments, as well as two segments of basic plus hydrophobic amino acids: residues 12-17 (pb1) and residues 26-33 (pb2) (Fig 1). Moreover, the C-terminally prenylated Rheb and RhoA lack secondary structure whereas the N-terminally myristoylated DNTE has a significant helical content (discussed above). Therefore, DNTE has the potential to interact with lipids via a combination of hydrophobic residues, an amphipathic helix, and two PBDs. Fig 6 A&B show that this is indeed the case. Almost all residues in the np segment, which are mostly located on the C-terminal helix, are deeply inserted into the bilayer and interact with lipid acyl chains. These include Leu 18, Leu 22, Ile 24 and Leu 25. Similarly, both the hydrophobic (e.g., Leu 13 and Leu 16) and basic (Lys 14, Arg 15 and Arg 17) residues in pb1 interact with lipids, with the Arg and Lys snorkeling to engage in both vdW and electrostatic interactions. The N-terminal half of pb2 residues including those on the helix, such as Arg 26, Phe 28 and Lys 29, engage lipids in a similar manner but the last few residues remain flexible and occasionally solvent exposed. The polar segment p forms a hook-like structure so that most side chains remain solvent exposed throughout the simulation while the myristate and Phe 5 insert deep into the bilayer hydrophobic core. These interactions are consistent with experimental observations. For example, simultaneously replacing 5 basic amino acids to Gln in pb1 and pb2 (Lys 11, Lys 14, Arg 15, Arg 17 and Arg 26) eliminated DNTE’s ability to bind anionic lipids but did not affect helicity^25^, and mutating the hydrophobic Leu and Ile residues in the middle of the peptide (in pb1 and np) eliminated both helicity and lipid binding ability^25^. Taking these together, we conclude that the basic residues of DNTE contribute to membrane binding whereas the non-polar amino acids in the center of the peptide have roles both in structural integrity and membrane binding.

**Figure 6:**
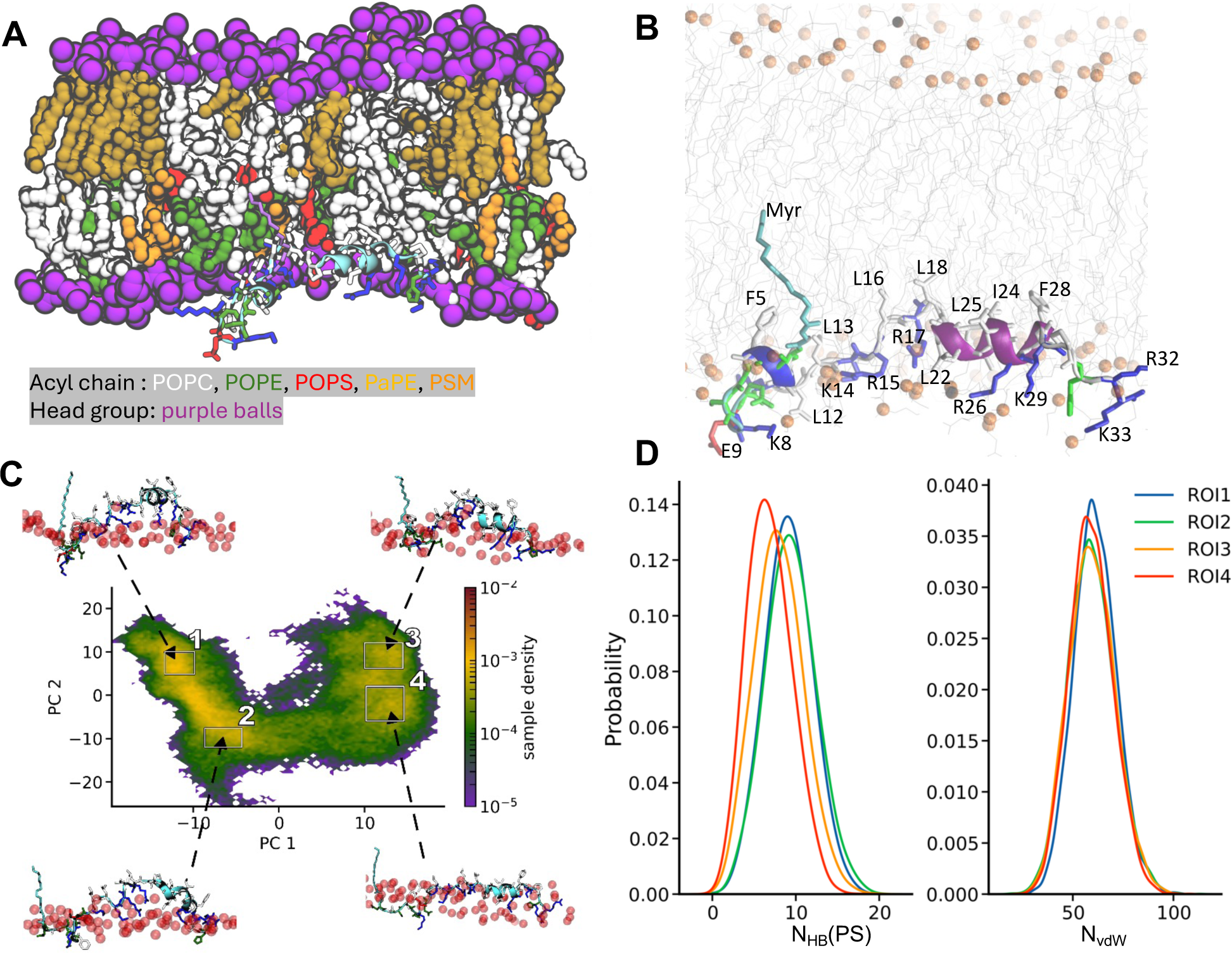
(**A**) A snapshot at the end of the DNTE simulation illustrating the organization of the peptide on the host monolayer with residues colored as in Fig 1D, except for the myristoylated Gly which is now in magenta. Lipid atoms including those of POPS within 10 Å of protein heavy atoms as well water and ions are omitted for clarity. (**B**) A close-up view of the bilayer insertion depth and lipid interaction of DNTE. Lipids are in grey lines with prosperous atoms shown as orange balls. DNTE is shown in cartoon representation with side chains colored in red (acidic), blue (basic), white (non-polar), and green (polar). (**C**) Distribution of simulated conformers along the top two principal components, PC1 and PC2, based on pair-wise Cα atom distances excluding adjacent residues. Example structures for the high-density regions labeled 1-4 are shown (arrows), highlighting the shape and insertion depth of DNTE relative to the phosphate groups of the host monolayer (red spheres). Side chains are shown in licorice with residues colored as in Fig 1. (**D**) Frequency of DNTE-PS hydrogen bonds (N_HB_, left) and van der Waals contacts (N_vdW_) of DNTE non-polar side chains and lipid acyl chains. Calculations were done using conformers from the regions of interest (ROI) marked with black squares in panel **B**.

### Electrostatic interactions with anionic lipids stabilize semi-ordered structures of RhoA and DNTE at membrane surfaces

We previously observed that extended and curled or semi-ordered conformations of KRAS^19^, RhoA and other PBD-containing lipid anchors of small GTPases^20,49^ interact differently with PS lipids. In the current work, we wondered if the frequently sampled RhoA conformations differ from the rest in their electrostatic interactions with lipids. To test this, we pooled together all meta-stable conformers (high density regions in Fig 5A) and compared their average N_HB_ with PS, PC and PE to that observed in all simulated conformers and in a pool of non-metastable conformers. We found that the number of HB contacts of RhoA with POPC is similar in the different pools of conformers but the meta-stable conformations have a higher total N_HB_ as well as PS- and PE-specific N_HB_s (Fig 5B). This shows that increased levels of hydrogen bonding interactions with anionic lipids stabilizes distinct conformational sub-states including semi-ordered conformations of RhoA.

To test if this observation on RhoA extends to the much longer and more complex IDR of DiRas3, we first performed principal component (PC) analysis over the entire trajectory based on pair-wise distances between non-adjacent Cα atoms (see methods). The goal was to categorize the simulated conformers into a manageable set of sub-ensembles. Fig 6C shows the resulting distribution of DNTE conformers along the top 2 PCs, which together covered over 55 % of the variance. We observe two well separated ensembles of conformers along PC1, each of which further divided in to at least two sub-ensembles along PC2. The snapshots taken from the center of the four high density regions or sub-ensembles (labeled 1-4 in Fig 6C) demonstrate a clear relationship between backbone conformation and interaction including membrane insertion depth (Fig 6C). In sub-ensembles 1 and 2, the peptide is bent at the center and inserts deep into the bilayer, with the two sub-ensembles differing primarily in the extent of bilayer penetration. The backbone is straighter in sub-ensembles 3 and 4 and lies mostly flat at the surface close to the phosphate group region. We then tested if these sub-ensembles differ in interactions with lipids by comparing the number of hydrogen bonds with PS (N_HB_(PS)) and the number of vdW contacts with lipid acyl chain carbons (N_vdW_). There is a clear correlation between backbone conformation and thus membrane insertion depth and N_HB_(PS), but not N_vdW_ (Fig 6D): deeply inserted conformations interact with PS phosphate oxygens less frequently.

Taken together, these results suggest that absence of preferred backbone conformations in Rheb can be explained by the lack of significant electrostatic interactions with lipids. The correlation between conformational preferences of RhoA and DNTE and side chain-dominated interactions with inner leaflet PM lipids supports previous observations on ensemble-dependent lipid sorting and a key role of the PBD in this process^24,25,49^.

### Protein-membrane electrostatic interactions alter local lipid composition

Because previous studies have indicated that the PBD of small GTPase membrane anchors induces clustering of anionic membrane lipids^14,15,19,20,49^, we decided to look at lipid clustering around the peptides simulated in this work. To do this, we calculated the 2D particle density of lipids around each peptide’s center of mass. We found significant clustering of anionic PS lipids around the PBD-containing DNTE and RhoA, but not the PBD-lacking Rheb, with the density of other lipids around DNTE being much lower (Fig 7). This result is consistent with the reduced lateral mobility of lipids in the peptide-containing leaflet (Fig 2B) and concordant with our previous observations in the KRAS membrane anchor^14^. Although there appears to be high density areas of SAPI around the Rheb anchor, the radial pair distribution (g(r)) plots in Fig S7A show that there is little interaction between the peptide and the anionic SAPI. To resolve this apparent contradiction, we measured the g(r) of each lipid type against itself. We found that the g(r) values for SAPI self-interaction were much higher than any other lipid type (Fig 7SB). Combined with experimental studies showing that SAPI forms microdomains^58,59^, we conclude that SAPI clustering is induced by self-interaction rather than interaction with Rheb. In contrast, there was little PS self-interaction in either of our PS-containing systems, confirming that the clustering of PS lipids we observe in Fig 7 is caused by selective interaction and sorting of this lipids by RhoA and DNTE.

**Figure 7:**
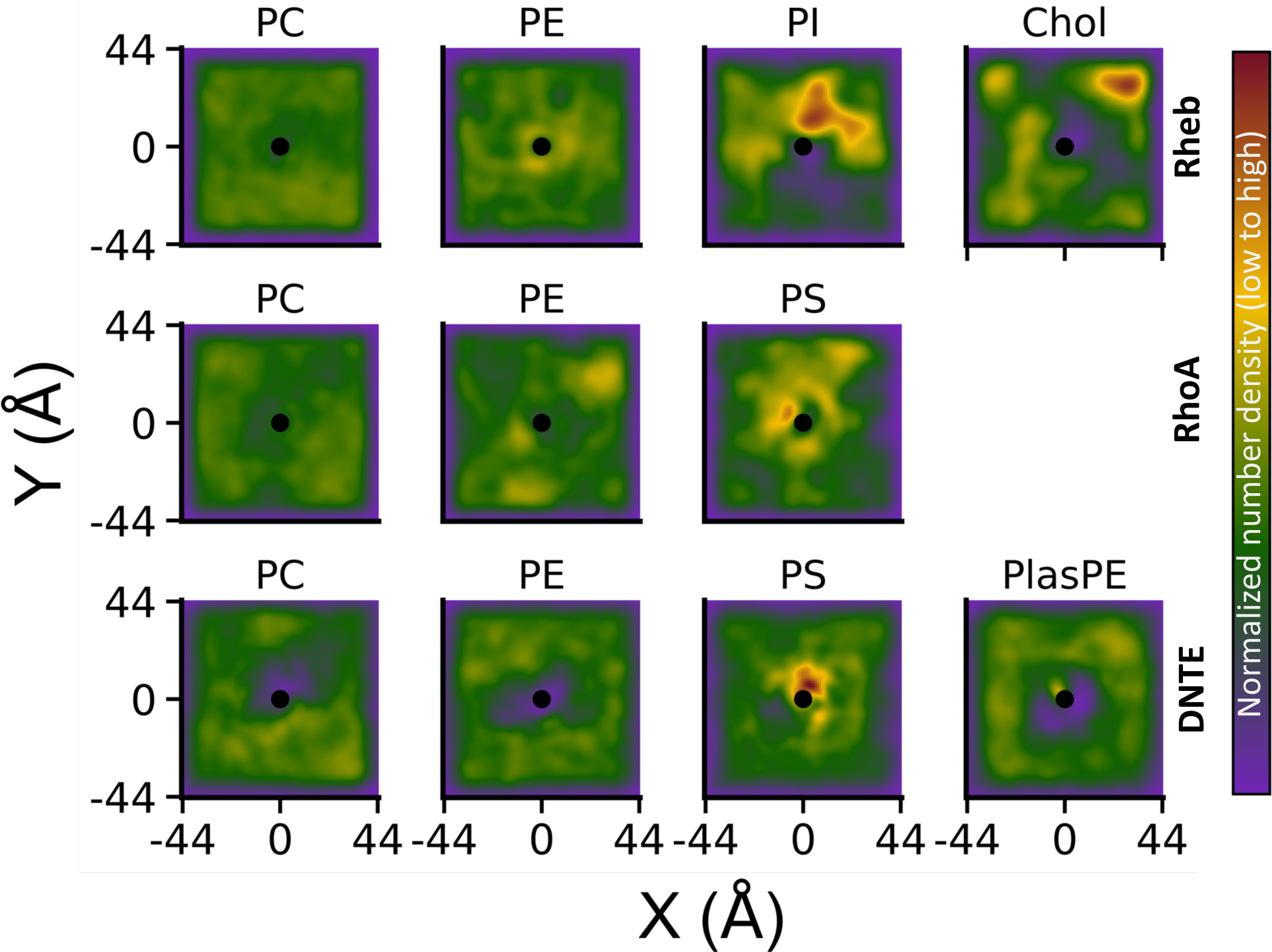
2D particle density of lipids around each lipidated IDR during the three simulations (see methods).

### Conclusion and implications for function

The three membrane anchors studied in this work represent distinct classes of lipidated IDRs in terms of both sequence composition and lipid modification (Fig 1). The C-terminally prenylated Rheb and RhoA are both 11-residues long, highly polar, and lack a secondary structure while the N-terminally myristoylated DNTE is much longer, more complex in sequence composition, and is partially helical. In addition, the small GTPases from which the three membrane anchors were derived diverge in their preference for cellular membranes, with Rheb primarily residing in ER membranes, RhoA in the PM, and DiRas3 more widely distributed but with preference for the PM. Therefore, we simulated each peptide in a different model membrane: Rheb in a bilayer of lipid composition approximating that of the ER, RhoA in a simplified PM model membrane, and DNTE in a complex bilayer following exploratory simulations in simpler bilayers. Results from the simulations indicate that the three lipidated IDRs display a wide range of membrane organization profile and conformational dynamics, and sampling of semi-stable conformational sub-states is a consequence of the combined effects of target membrane lipid composition and IDR sequence. Specifically, the PBD of RhoA and DNTE stabilizes specific conformational sub-states by interacting with anionic lipids, in agreement with observations in the KRAS membrane anchor^19^. In contrast, the IDR of Rheb lacks not only a PBD but also a net charge, and therefore only weakly interacts with PE but not with the anionic SAPI. The plasma membrane is highly enriched with anionic lipids, while the ER carries a much more neutral charge. This contrast highlights the importance of complementarity between a lipidated IDR sequence and lipid composition of its target membrane. The high density of polar residues in Rheb may also reduce its membrane affinity, and the lack of favorable electrostatic or hydrophobic interactions prevent formation of meta-stable conformational sub-states. These insights may provide a molecular basis for how lipidated membrane anchor sequence in tandem with target membrane lipid composition work together to determine membrane orientation dynamics of small GTPases.

Interaction with negatively charged phospholipids is believed to be an important mechanism by which KRAS forms nanoclusters on the PM to enhance signaling efficacy^60^. Besides KRAS, there is evidence that other small GTPases, such as Rac1 and Cdc42, also form nanoclusters that may enhance their signaling potential^9,39–41^. Although there is no clear evidence for RhoA or DNTE forming nanoclusters^61^, our findings that their membrane anchors preferentially interact with and segregate anionic lipids imply that these proteins may also be able to form signaling nanoclusters through a similar lipid sorting mechanism. Moreover, DNTE segregated PS lipids to such a degree that other lipid species were far less present around the peptide (Fig 7). This result corroborates experimental studies that have shown that DNTE interacts strongly with anionic PIP lipids and, albeit with less affinity, with PS lipids^25^. While DiRas3 has been shown to prevent formation of KRAS nanoclusters, possibly through competitive dimerization with KRAS^31^, our observations on DNTE’s ability to displace other lipid species in favor of PS suggests that competitive interactions with anionic phospholipids may also contribute to the disruption of KRAS nanoclusters whose stability has been shown to be PS dependent^60,62^.

## Acknowledgements

This work was supported in part by the National Institutes of Health Institute of General Medicine grant R01GM144836 to AAG. CMH was supported by an F32 Fellowship grant # T32GM139801. Computational resources have been provided by the Texas Advanced Computing Center (TACC) and Anton 2. Anton 2 computer time was provided by the Pittsburgh Supercomputing Center (PSC) through Grant R01GM116961 from the National Institutes of Health. The Anton 2 machine at PSC was generously made available by D.E. Shaw Research. We thank Violeta Burns Casamayor for running the very early exploratory simulations.

## Author contributions

A.A.G. conceived and designed the project; C.M.H. performed the simulations; C.M.H. and A.A.G. analyzed the data and wrote the paper.

## Competing interests

The authors declare no competing interests.

## Notes

### Competing Interest Statement

The authors have declared no competing interest.

## References

(1) Oldfield, C. J.; Dunker, A. K. Intrinsically Disordered Proteins and Intrinsically Disordered Protein Regions. Annu. Rev. Biochem. 2014, 83, 553–584. 10.1146/annurev-biochem-072711-164947.

(2) Granata, D.; Baftizadeh, F.; Habchi, J.; Galvagnion, C.; De Simone, A.; Camilloni, C.; Laio, A.; Vendruscolo, M. The Inverted Free Energy Landscape of an Intrinsically Disordered Peptide by Simulations and Experiments. Sci. Rep. 2015, 5, 15449. 10.1038/srep15449.

(3) Wright, P. E.; Dyson, H. J. Intrinsically Disordered Proteins in Cellular Signaling and Regulation. Nat. Rev. Mol. Cell Biol. 2015, 16 (1), 18–29. 10.1038/nrm3920.

(4) Iakoucheva, L. M.; Brown, C. J.; Lawson, J. D.; Obradović, Z.; Dunker, A. K. Intrinsic Disorder in Cell-Signaling and Cancer-Associated Proteins. J. Mol. Biol. 2002, 323 (3), 573–584. 10.1016/s0022-2836(02)00969-5.

(5) Braicu, C.; Buse, M.; Busuioc, C.; Drula, R.; Gulei, D.; Raduly, L.; Rusu, A.; Irimie, A.; Atanasov, A. G.; Slaby, O.; Ionescu, C.; Berindan-Neagoe, I. A Comprehensive Review on MAPK: A Promising Therapeutic Target in Cancer. Cancers 2019, 11 (10), 1618. 10.3390/cancers11101618.

(6) Goitre, L.; Trapani, E.; Trabalzini, L.; Retta, S. F. The Ras Superfamily of Small GTPases: The Unlocked Secrets. In Ras Signaling: Methods and Protocols; Trabalzini, L., Retta, S. F., Eds.; Methods in Molecular Biology; Humana Press: Totowa, NJ, 2014; pp 1–18. 10.1007/978-1-62703-791-4_1.

(7) Wennerberg, K.; Rossman, K. L.; Der, C. J. The Ras Superfamily at a Glance. J. Cell Sci. 2005, 118 (5), 843–846. 10.1242/jcs.01660.

(8) Macara, I. G.; Lounsbury, K. M.; Richards, S. A.; McKiernan, C.; Bar-Sagi, D. The Ras Superfamily of GTPases. FASEB J. Off. Publ. Fed. Am. Soc. Exp. Biol. 1996, 10 (5), 625– 630. 10.1096/fasebj.10.5.8621061.

(9) Cox, A. D.; Der, C. J. Ras Family Signaling: Therapeutic Targeting. Cancer Biol. Ther. 2002, 1 (6), 599–606. 10.4161/cbt.306.

(10) Qu, L.; Pan, C.; He, S.-M.; Lang, B.; Gao, G.-D.; Wang, X.-L.; Wang, Y. The Ras Superfamily of Small GTPases in Non-Neoplastic Cerebral Diseases. Front. Mol. Neurosci. 2019, 12, 121. 10.3389/fnmol.2019.00121.

(11) Vetter, I. R.; Wittinghofer, A. The Guanine Nucleotide-Binding Switch in Three Dimensions. Science 2001, 294 (5545), 1299–1304. 10.1126/science.1062023.

(12) Hancock, J. F.; Magee, A. I.; Childs, J. E.; Marshall, C. J. All Ras Proteins Are Polyisoprenylated but Only Some Are Palmitoylated. Cell 1989, 57 (7), 1167–1177. 10.1016/0092-8674(89)90054-8.

(13) Hancock, J. F. Ras Proteins: Different Signals from Different Locations. Nat. Rev. Mol. Cell Biol. 2003, 4 (5), 373–385. 10.1038/nrm1105.

(14) Zhou, Y.; Prakash, P.; Liang, H.; Cho, K.-J.; Gorfe, A. A.; Hancock, J. F. Lipid-Sorting Specificity Encoded in K-Ras Membrane Anchor Regulates Signal Output. Cell 2017, 168 (1), 239–251.e16. 10.1016/j.cell.2016.11.059.

(15) Zhou, Y.; Prakash, P. S.; Liang, H.; Gorfe, A. A.; Hancock, J. F. The KRAS and Other Prenylated Polybasic Domain Membrane Anchors Recognize Phosphatidylserine Acyl Chain Structure. Proc. Natl. Acad. Sci. 2021, 118 (6), e2014605118. 10.1073/pnas.2014605118.

(16) Prakash, P.; Zhou, Y.; Liang, H.; Hancock, J. F.; Gorfe, A. A. Oncogenic K-Ras Binds to an Anionic Membrane in Two Distinct Orientations: A Molecular Dynamics Analysis. Biophys. J. 2016, 110 (5), 1125–1138. 10.1016/j.bpj.2016.01.019.

(17) Prakash, P.; Gorfe, A. A. Determinants of Membrane Orientation Dynamics in Lipid-Modified Small GTPases. JACS Au 2022, 2 (1), 128–135. 10.1021/jacsau.1c00426.

(18) Prakash, P.; Gorfe, A. A. Membrane Orientation Dynamics of Lipid-Modified Small GTPases. Small GTPases 2016, 8 (3), 129–138. 10.1080/21541248.2016.1211067.

(19) Janosi, L.; Gorfe, A. A. Segregation of Negatively Charged Phospholipids by the Polycationic and Farnesylated Membrane Anchor of Kras. Biophys. J. 2010, 99 (11), 3666–3674. 10.1016/j.bpj.2010.10.031.

(20) Araya, M. K.; Gorfe, A. A. Phosphatidylserine and Phosphatidylethanolamine Asymmetry Have a Negligible Effect on the Global Structure, Dynamics, and Interactions of the KRAS Lipid Anchor. J. Phys. Chem. B 2022, 126 (24), 4491–4500. 10.1021/acs.jpcb.2c01253.

(21) Angarola, B.; Ferguson, S. M. Weak Membrane Interactions Allow Rheb to Activate mTORC1 Signaling without Major Lysosome Enrichment. Mol. Biol. Cell 2019, 30 (22), 2750–2760. 10.1091/mbc.E19-03-0146.

(22) Saucedo, L. J.; Gao, X.; Chiarelli, D. A.; Li, L.; Pan, D.; Edgar, B. A. Rheb Promotes Cell Growth as a Component of the Insulin/TOR Signalling Network. Nat. Cell Biol. 2003, 5 (6), 566–571. 10.1038/ncb996.

(23) Jaffe, A. B.; Hall, A. Rho GTPases: Biochemistry and Biology. Annu. Rev. Cell Dev. Biol. 2005, 21, 247–269. 10.1146/annurev.cellbio.21.020604.150721.

(24) Yu, Y.; Xu, F.; Peng, H.; Fang, X.; Zhao, S.; Li, Y.; Cuevas, B.; Kuo, W. L.; Gray, J. W.; Siciliano, M.; Mills, G. B.; Bast, R. C. NOEY2 (ARHI), an Imprinted Putative Tumor Suppressor Gene in Ovarian and Breast Carcinomas. Proc. Natl. Acad. Sci. U. S. A. 1999, 96 (1), 214–219. 10.1073/pnas.96.1.214.

(25) Liang, X.; Jung, S. Y.; Fong, L. W.; Bildik, G.; Gray, J. P.; Mao, W.; Zhang, S.; Millward, S. W.; Gorfe, A. A.; Zhou, Y.; Lu, Z.; Bast, R. C. Membrane Anchoring of the DIRAS3 N-Terminal Extension Permits Tumor Suppressor Function. iScience 2023, 26 (11), 108151. 10.1016/j.isci.2023.108151.

(26) Mark, P.; Nilsson, L. Structure and Dynamics of the TIP3P, SPC, and SPC/E Water Models at 298 K. J. Phys. Chem. A 2001, 105 (43), 9954–9960. 10.1021/jp003020w.

(27) Jo, S.; Kim, T.; Iyer, V. G.; Im, W. CHARMM-GUI: A Web-Based Graphical User Interface for CHARMM. J. Comput. Chem. 2008, 29 (11), 1859–1865. 10.1002/jcc.20945.

(28) Jo, S.; Lim, J. B.; Klauda, J. B.; Im, W. CHARMM-GUI Membrane Builder for Mixed Bilayers and Its Application to Yeast Membranes. Biophys. J. 2009, 97 (1), 50–58. 10.1016/j.bpj.2009.04.013.

(29) van Meer, G.; Voelker, D. R.; Feigenson, G. W. Membrane Lipids: Where They Are and How They Behave. Nat. Rev. Mol. Cell Biol. 2008, 9 (2), 112–124. 10.1038/nrm2330.

(30) Prakash, P.; Litwin, D.; Liang, H.; Sarkar-Banerjee, S.; Dolino, D.; Zhou, Y.; Hancock, J. F.; Jayaraman, V.; Gorfe, A. A. Dynamics of Membrane-Bound G12V-KRAS from Simulations and Single-Molecule FRET in Native Nanodiscs. Biophys. J. 2019, 116 (2), 179–183. 10.1016/j.bpj.2018.12.011.

(31) Sutton, M. N.; Lu, Z.; Li, Y.-C.; Zhou, Y.; Huang, T.; Reger, A. S.; Hurwitz, A. M.; Palzkill, T.; Logsdon, C.; Liang, X.; Gray, J. W.; Nan, X.; Hancock, J.; Wahl, G. M.; Bast, R. C. DIRAS3 (ARHI) Blocks RAS/MAPK Signaling by Binding Directly to RAS and Disrupting RAS Clusters. Cell Rep. 2019, 29 (11), 3448–3459.e6. 10.1016/j.celrep.2019.11.045.

(32) Lorent, J. H.; Levental, K. R.; Ganesan, L.; Rivera-Longsworth, G.; Sezgin, E.; Doktorova, M.; Lyman, E.; Levental, I. Plasma Membranes Are Asymmetric in Lipid Unsaturation, Packing and Protein Shape. Nat. Chem. Biol. 2020, 16 (6), 644–652. 10.1038/s41589-020-0529-6.

(33) Huang, J.; MacKerell, A. D. CHARMM36 All-Atom Additive Protein Force Field: Validation Based on Comparison to NMR Data. J. Comput. Chem. 2013, 34 (25), 2135–2145. 10.1002/jcc.23354.

(34) Phillips, J. C.; Hardy, D. J.; Maia, J. D. C.; Stone, J. E.; Ribeiro, J. V.; Bernardi, R. C.; Buch, R.; Fiorin, G.; Hénin, J.; Jiang, W.; McGreevy, R.; Melo, M. C. R.; Radak, B. K.; Skeel, R. D.; Singharoy, A.; Wang, Y.; Roux, B.; Aksimentiev, A.; Luthey-Schulten, Z.; Kalé, L. V.; Schulten, K.; Chipot, C.; Tajkhorshid, E. Scalable Molecular Dynamics on CPU and GPU Architectures with NAMD. J. Chem. Phys. 2020, 153 (4), 044130. 10.1063/5.0014475.

(35) Abraham, M. J.; Murtola, T.; Schulz, R.; Páll, S.; Smith, J. C.; Hess, B.; Lindahl, E. GROMACS: High Performance Molecular Simulations through Multi-Level Parallelism from Laptops to Supercomputers. SoftwareX 2015, 1–2, 19–25. 10.1016/j.softx.2015.06.001.

(36) Darden, T.; York, D.; Pedersen, L. Particle Mesh Ewald: An N⋅log(N) Method for Ewald Sums in Large Systems. J. Chem. Phys. 1993, 98 (12), 10089–10092. 10.1063/1.464397.

(37) Elber, R.; Ruymgaart, A. P.; Hess, B. SHAKE Parallelization. Eur. Phys. J. Spec. Top. 2011, 200 (1), 211–223. 10.1140/epjst/e2011-01525-9.

(38) Hess, B.; Bekker, H.; Berendsen, H. J. C.; Fraaije, J. G. E. M. LINCS: A Linear Constraint Solver for Molecular Simulations. J. Comput. Chem. 1997, 18 (12), 1463–1472. 10.1002/(SICI)1096-987X(199709)18:12<1463::AID-JCC4>3.0.CO;2-H.

(39) Bussi, G.; Donadio, D.; Parrinello, M. Canonical Sampling through Velocity Rescaling. J. Chem. Phys. 2007, 126 (1), 014101. 10.1063/1.2408420.

(40) Parrinello, M.; Rahman, A. Polymorphic Transitions in Single Crystals: A New Molecular Dynamics Method. J. Appl. Phys. 1981, 52 (12), 7182–7190. 10.1063/1.328693.

(41) Shaw, D. E.; Grossman, J. P.; Bank, J. A.; Batson, B.; Butts, J. A.; Chao, J. C.; Deneroff, M. M.; Dror, R. O.; Even, A.; Fenton, C. H.; Forte, A.; Gagliardo, J.; Gill, G.; Greskamp, B.; Ho, C. R.; Ierardi, D. J.; Iserovich, L.; Kuskin, J. S.; Larson, R. H.; Layman, T.; Lee, L.-S.; Lerer, A. K.; Li, C.; Killebrew, D.; Mackenzie, K. M.; Mok, S. Y.-H.; Moraes, M. A.; Mueller, R.; Nociolo, L. J.; Peticolas, J. L.; Quan, T.; Ramot, D.; Salmon, J. K.; Scarpazza, D. P.; Schafer, U. B.; Siddique, N.; Snyder, C. W.; Spengler, J.; Tang, P. T. P.; Theobald, M.; Toma, H.; Towles, B.; Vitale, B.; Wang, S. C.; Young, C. Anton 2: Raising the Bar for Performance and Programmability in a Special-Purpose Molecular Dynamics Supercomputer. In SC ‘14: Proceedings of the International Conference for High Performance Computing, Networking, Storage and Analysis; 2014; pp 41–53. 10.1109/SC.2014.9.

(42) Gowers, R. J.; Linke, M.; Barnoud, J.; Reddy, T. J. E.; Melo, M. N.; Seyler, S. L.; Domański, J.; Dotson, D. L.; Buchoux, S.; Kenney, I. M.; Beckstein, O. MDAnalysis: A Python Package for the Rapid Analysis of Molecular Dynamics Simulations. Proc. 15th Python Sci. Conf. 2016, 98–105. 10.25080/Majora-629e541a-00e.

(43) Humphrey, W.; Dalke, A.; Schulten, K. VMD: Visual Molecular Dynamics. J. Mol. Graph. 1996, 14 (1), 33–38, 27–28. 10.1016/0263-7855(96)00018-5.

(44) Virtanen, P.; Gommers, R.; Oliphant, T. E.; Haberland, M.; Reddy, T.; Cournapeau, D.; Burovski, E.; Peterson, P.; Weckesser, W.; Bright, J.; van der Walt, S. J.; Brett, M.; Wilson, J.; Millman, K. J.; Mayorov, N.; Nelson, A. R. J.; Jones, E.; Kern, R.; Larson, E.; Carey, C. J.; Polat, İ.; Feng, Y.; Moore, E. W.; VanderPlas, J.; Laxalde, D.; Perktold, J.; Cimrman, R.; Henriksen, I.; Quintero, E. A.; Harris, C. R.; Archibald, A. M.; Ribeiro, A. H.; Pedregosa, F.; van Mulbregt, P. SciPy 1.0: Fundamental Algorithms for Scientific Computing in Python. Nat. Methods 2020, 17 (3), 261–272. 10.1038/s41592-019-0686-2.

(45) Smith, P.; Lorenz, C. D. LiPyphilic: A Python Toolkit for the Analysis of Lipid Membrane Simulations. J. Chem. Theory Comput. 2021, 17 (9), 5907–5919. 10.1021/acs.jctc.1c00447.

(46) Scherer, M. K.; Trendelkamp-Schroer, B.; Paul, F.; Pérez-Hernández, G.; Hoffmann, M.; Plattner, N.; Wehmeyer, C.; Prinz, J.-H.; Noé, F. PyEMMA 2: A Software Package for Estimation, Validation, and Analysis of Markov Models. J. Chem. Theory Comput. 2015, 11 (11), 5525–5542. 10.1021/acs.jctc.5b00743.

(47) Kabsch, W.; Sander, C. Dictionary of Protein Secondary Structure: Pattern Recognition of Hydrogen-Bonded and Geometrical Features. Biopolymers 1983, 22 (12), 2577–2637. 10.1002/bip.360221211.

(48) McGibbon, R. T.; Beauchamp, K. A.; Harrigan, M. P.; Klein, C.; Swails, J. M.; Hernández, C. X.; Schwantes, C. R.; Wang, L.-P.; Lane, T. J.; Pande, V. S. MDTraj: A Modern Open Library for the Analysis of Molecular Dynamics Trajectories. Biophys. J. 2015, 109 (8), 1528–1532. 10.1016/j.bpj.2015.08.015.

(49) Araya, M. K.; Gorfe, A. A. Conformational Ensemble-Dependent Lipid Recognition and Segregation by Prenylated Intrinsically Disordered Regions in Small GTPases. Commun. Biol. 2023, 6, 1111. 10.1038/s42003-023-05487-6.

(50) Leftin, A.; Molugu, T. R.; Job, C.; Beyer, K.; Brown, M. F. Area per Lipid and Cholesterol Interactions in Membranes from Separated Local-Field 13C NMR Spectroscopy. Biophys. J. 2014, 107 (10), 2274–2286. 10.1016/j.bpj.2014.07.044.

(51) Ferreira, T. M.; Coreta-Gomes, F.; Ollila, O. H. S.; Moreno, M. J.; Vaz, W. L. C.; Topgaard, D. Cholesterol and POPC Segmental Order Parameters in Lipid Membranes: Solid State 1H-13C NMR and MD Simulation Studies. Phys. Chem. Chem. Phys. PCCP 2013, 15 (6), 1976–1989. 10.1039/c2cp42738a.

(52) Kučerka, N.; Nieh, M.-P.; Katsaras, J. Fluid Phase Lipid Areas and Bilayer Thicknesses of Commonly Used Phosphatidylcholines as a Function of Temperature. Biochim. Biophys. Acta BBA - Biomembr. 2011, 1808 (11), 2761–2771. 10.1016/j.bbamem.2011.07.022.

(53) Janosi, L.; Gorfe, A. A. Simulating POPC and POPC/POPG Bilayers: Conserved Packing and Altered Surface Reactivity. J. Chem. Theory Comput. 2010, 6 (10), 3267–3273. 10.1021/ct100381g.

(54) Filippov, A.; Orädd, G.; Lindblom, G. The Effect of Cholesterol on the Lateral Diffusion of Phospholipids in Oriented Bilayers. Biophys. J. 2003, 84 (5), 3079–3086.

(55) Drin, G.; Casella, J.-F.; Gautier, R.; Boehmer, T.; Schwartz, T. U.; Antonny, B. A General Amphipathic α-Helical Motif for Sensing Membrane Curvature. Nat. Struct. Mol. Biol. 2007, 14 (2), 138–146. 10.1038/nsmb1194.

(56) Cui, H.; Lyman, E.; Voth, G. A. Mechanism of Membrane Curvature Sensing by Amphipathic Helix Containing Proteins. Biophys. J. 2011, 100 (5), 1271–1279. 10.1016/j.bpj.2011.01.036.

(57) Giménez-Andrés, M.; Čopič, A.; Antonny, B. The Many Faces of Amphipathic Helices. Biomolecules 2018, 8 (3), 45. 10.3390/biom8030045.

(58) Motegi, T.; Takiguchi, K.; Tanaka-Takiguchi, Y.; Itoh, T.; Tero, R. Physical Properties and Reactivity of Microdomains in Phosphatidylinositol-Containing Supported Lipid Bilayer. Membranes 2021, 11 (5), 339. 10.3390/membranes11050339.

(59) Redfern, D. A.; Gericke, A. Domain Formation in Phosphatidylinositol Monophosphate/Phosphatidylcholine Mixed Vesicles. Biophys. J. 2004, 86 (5), 2980–2992. 10.1016/S0006-3495(04)74348-9.

(60) Zhou, Y.; Gorfe, A. A.; Hancock, J. F. RAS Nanoclusters Selectively Sort Distinct Lipid Headgroups and Acyl Chains. Front. Mol. Biosci. 2021, 8, 686338. 10.3389/fmolb.2021.686338.

(61) de Seze, J.; Gatin, J.; Coppey, M. RhoA Regulation in Space and Time. FEBS Lett. 2023, 597 (6), 836–849. 10.1002/1873-3468.14578.

(62) Zhou, Y.; Wong, C.-O.; Cho, K.; van der Hoeven, D.; Liang, H.; Thakur, D. P.; Luo, J.; Babic, M.; Zinsmaier, K. E.; Zhu, M. X.; Hu, H.; Venkatachalam, K.; Hancock, J. F. Membrane Potential Modulates Plasma Membrane Phospholipid Dynamics and K-Ras Signaling. Science 2015, 349 (6250), 873–876. 10.1126/science.aaa5619.

