## Supplemental text and figures for "Intrinsically disordered membrane anchors of Rheb, RhoA and DiRas3 small GTPases: Molecular dynamics, membrane organization, and interactions"

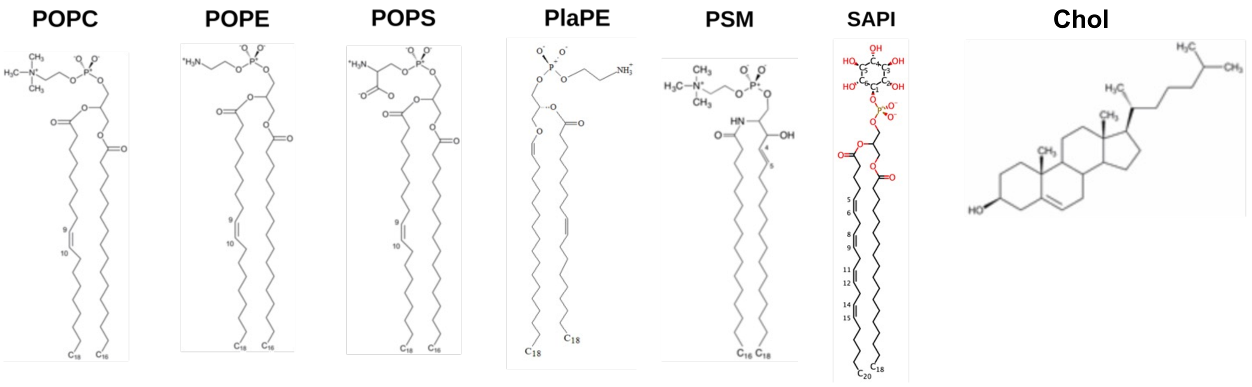


**Figure S1:** Molecular structure of lipids used in the current simulations (from https://charmm-gui.org/?doc=archive&lib=lipid). POPC = 16:1-18:0 1-palmitoyl-2-oleoyl-glycero-3-phosphocholine; POPE = 16:1-18:0 1-palmitoyl-2-oleoyl-glycero-3-phosphatidylethanolamine; POPS = 16:1-18:0 1-palmitoyl-2-oleoyl-glycero-3-phosphatidylserine; 18:1-18:1 plasmalogen phosphatidylethanolamine; SAPI = 18:0-20.4 phosphatidylinositol; Chol = Cholesterol.

***Generating initial structure of DNTE.*** We previously used online servers to predict multiple DNTE structures[1]. Most of these structures contained a helix-turn-helix motif and have a much higher helical content (~80%) than has been observed with CD in solution (~14%)[1]. Importantly, none of the structures appeared to be ideal for membrane binding. 3 μs-long MD simulations of three such models in a PC/PS bilayer did not fully resolve the issue. In two of the simulations, the helix-turn-helix structure persisted. In the third simulation, the overall helical content reduced to ~30% due to one of the helices melting. We therefore started a new simulation from the last snapshot of this run, attaching the peptide to the anionic monolayer of an asymmetric PC-PC/PS bilayer. A 2 μs MD run of this setup resulted in a stable system with convergent backbone root mean square derivation (RMSD) and secondary content (Fig S2). However, the amphipathic **α**-helix near the N-terminus (residues 8-14) adopted an orientation in which the hydrophobic face is exposed to solvent (see snapshot in Fig S2). This orientation is inconsistent with known patterns of amphipathic helix membrane binding[2–4]. To check if the helix might spontaneously flip to burry its hydrophobic face in the bilayer, we started a second simulation from the final snapshot of the 2 μs run using a more complex bilayer made up of a neutral POPC/PSM monolayer and an anionic POPC/POPS/POPE/PlaPE monolayer (see main text). We observed little change in the structure of the peptide or the membrane organization of the helix during a 10 μs simulation of this system (Fig S2), suggesting a large energy barrier to flipping the helix. Therefore, a third simulation was started from the last snapshot of this run after manually rotating the helix such that the hydrophobic face was embedded in the membrane and the polar face oriented to solvent. This setup allowed for non-polar residues to engage in vdW interactions with lipid acyl chains and basic residues to make both vdW contacts with acyl chains and electrostatic interactions with head groups. A 10 μs run of this system yielded reliable results, including DNTE structure and membrane binding profiles that are consistent with available experimental data for DNTE[1] and other amphipathic helices[2-4]. These results are discussed in the main manuscript.

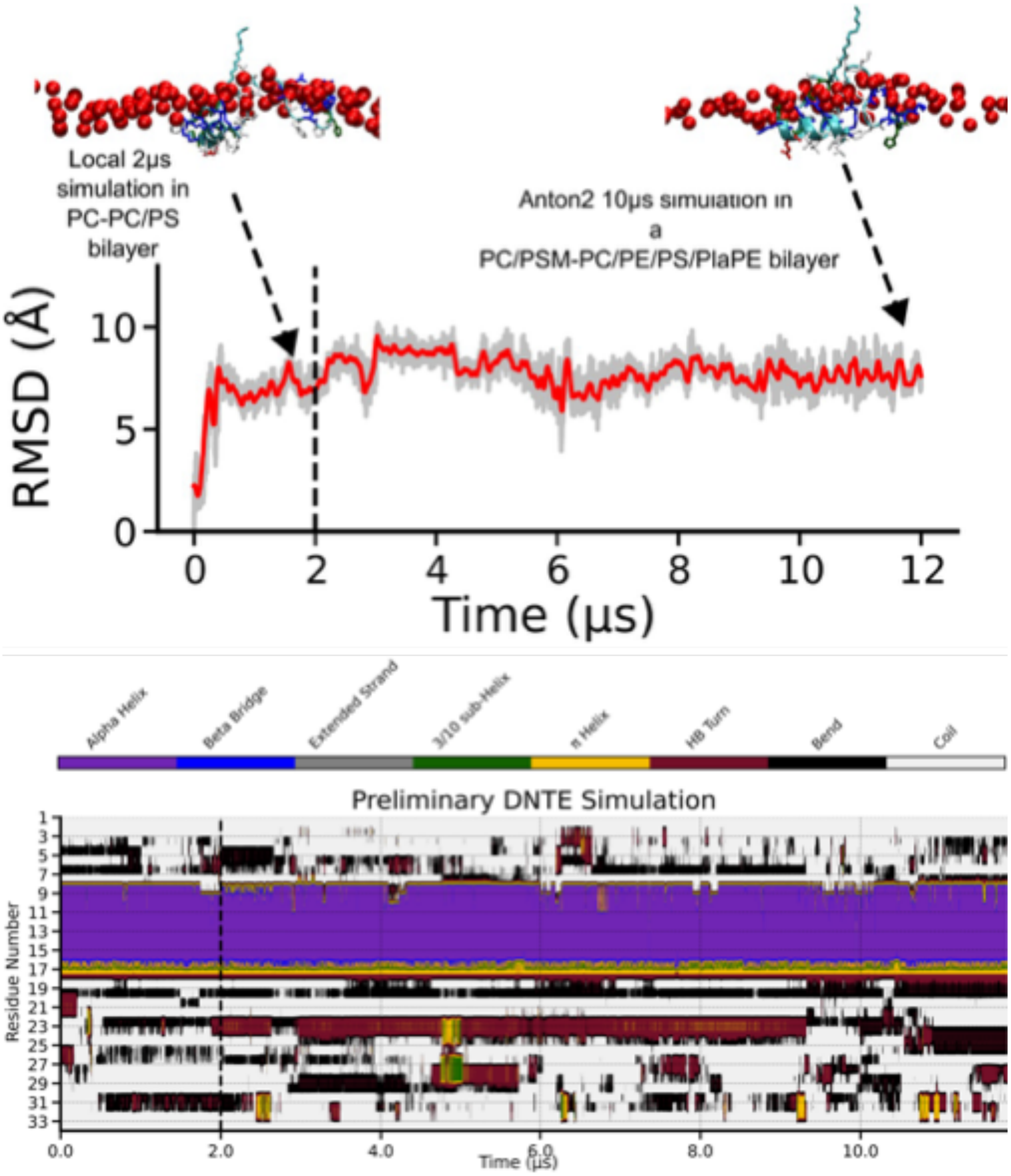


**Figure S2:** Time evolution of DNTE backbone RMSD (top) and secondary structure (bottom) during two exploratory MD simulations: a 2 μs run in a PC-PC/PS bilayer and a 10 μs run in a PC/PSM-PC/PE/PlaPE bilayer (demarcated by vertical dashed lines). The snapshots highlight the orientation of the amphipathic helix with its polar face toward solvent in both simulations. For clarity, only phosphates of the host monolayer are shown as red spheres. The helix is in cyan, basic residues in blue, non-polar in white, acidic in red and polar residues in green. Note the persistence of the N-terminal α-helix (residues 8-14) and the infrequent appearance of other secondary structures.

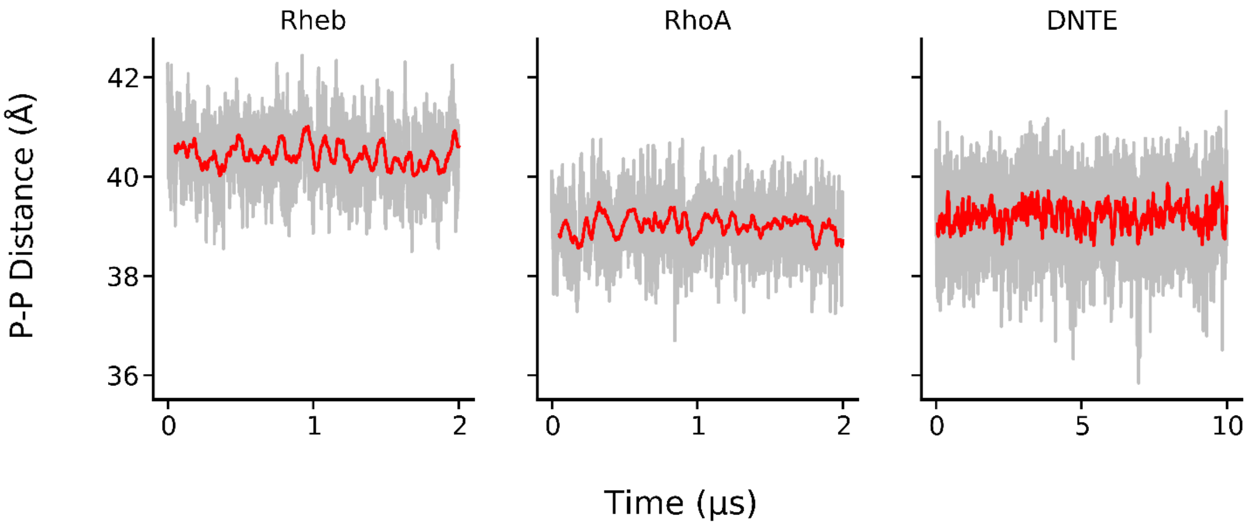


**Figure S3:** Time evolution of bilayer thickness (P-P distance) defined as the average z-distance between the centers-of-mass of the upper and lower leaflet phosphorous atoms in simulations Rheb, RhoA, and DNTE. The raw data is in light grey and 200 ns moving averages are in red.

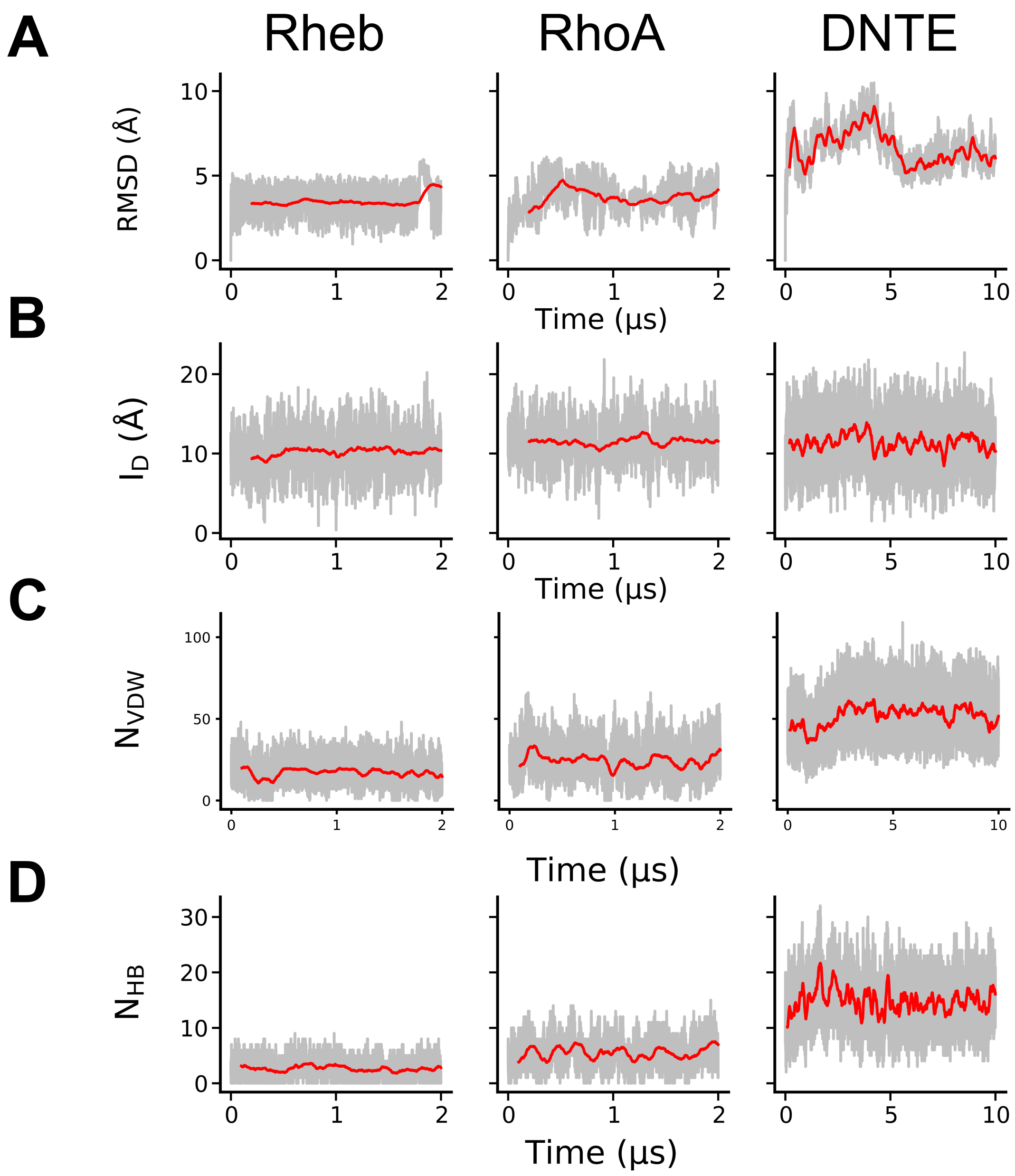


**Figure S4:** Time evolution of peptide backbone RMSD from the initial frames (**A**), insertion depth ($I_D$) of lipidated side chains **(B**), number of van der Waals contacts ($N_{vdW}$) (**C**), and number of hydrogen bonding interactions ($N_{HB}$) (**D**). The raw data is in light grey and 200 ns moving averages in red. See main text for definitions.

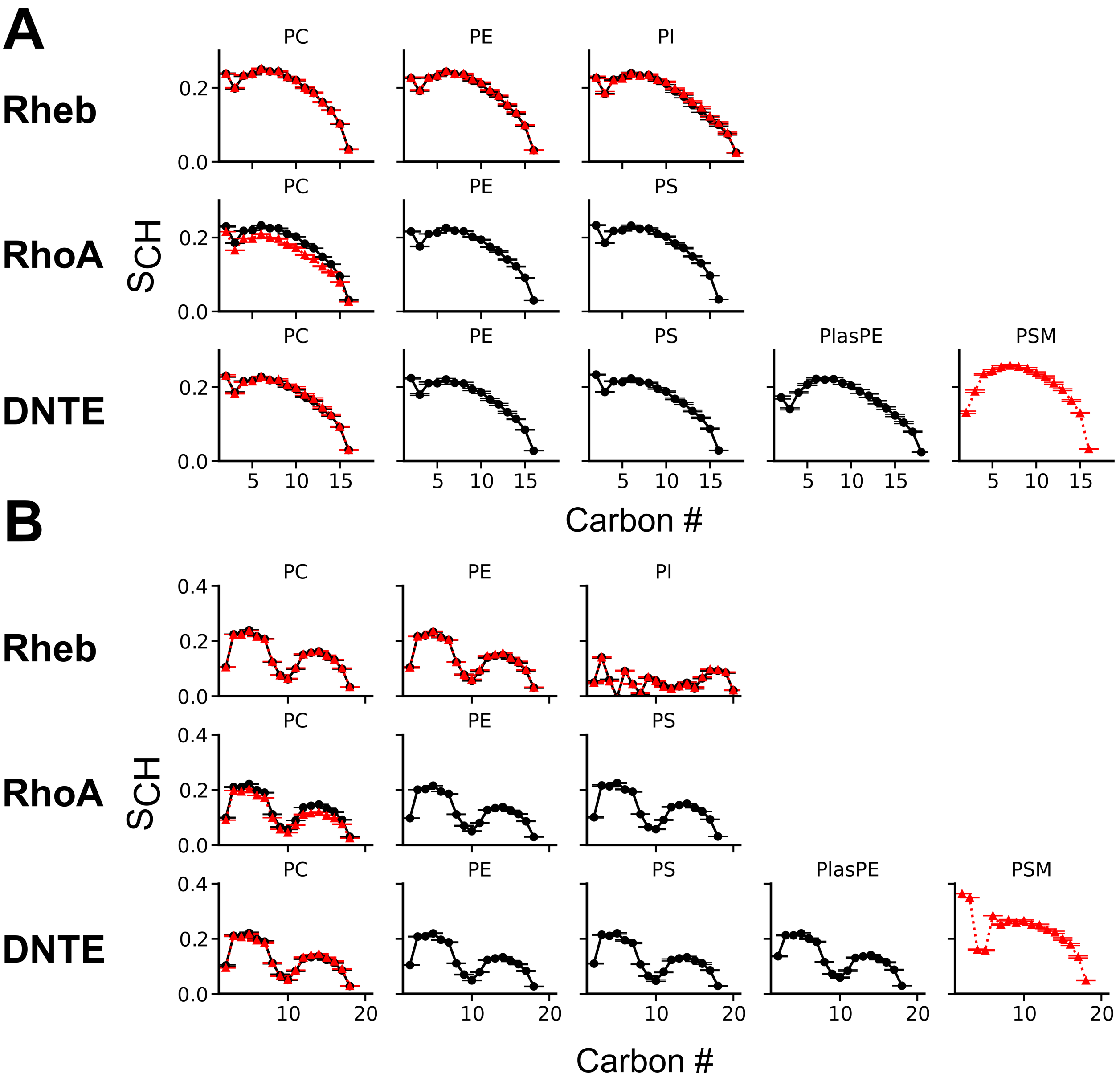


**Figure S5:** Sn-1 (**A**) and Sn-2 (**B**) acyl chain order parameters ($S_{CH}$) of indicated lipids in the upper (red) and lower (black) leaflets of bilayers in the Rheb, RhoA and DNTE simulations. See Table 1 of the main text for definition of lipid types.

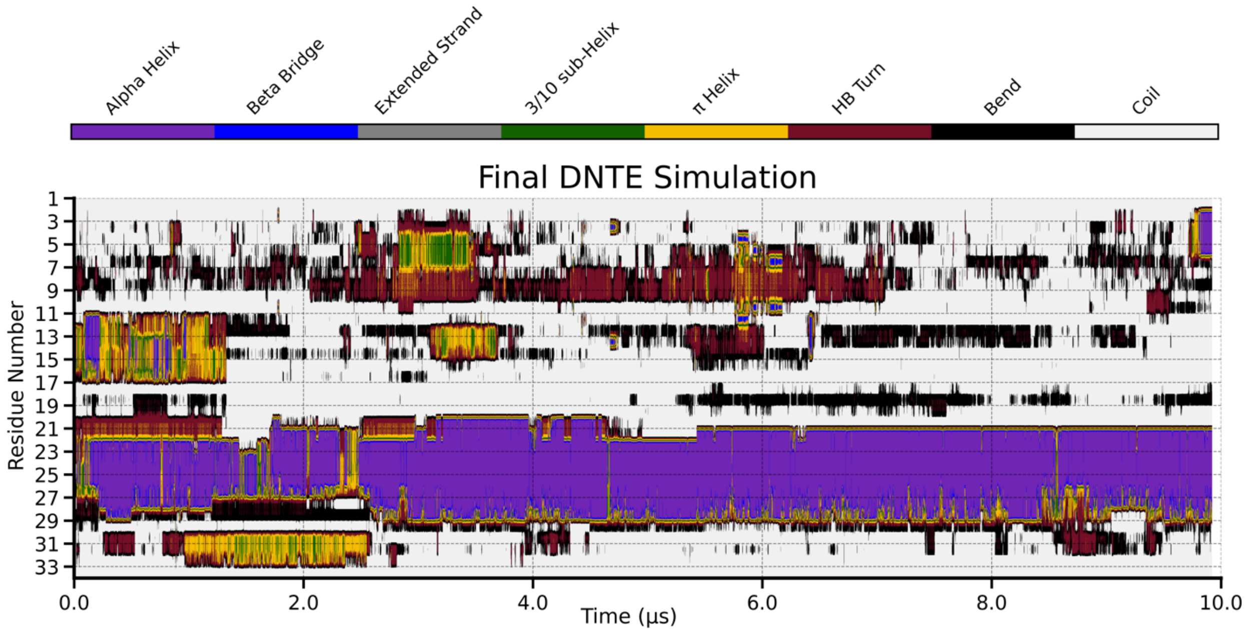


**Figure S6:** Evolution of DNTE secondary structure during the final simulation discussed in the main text.

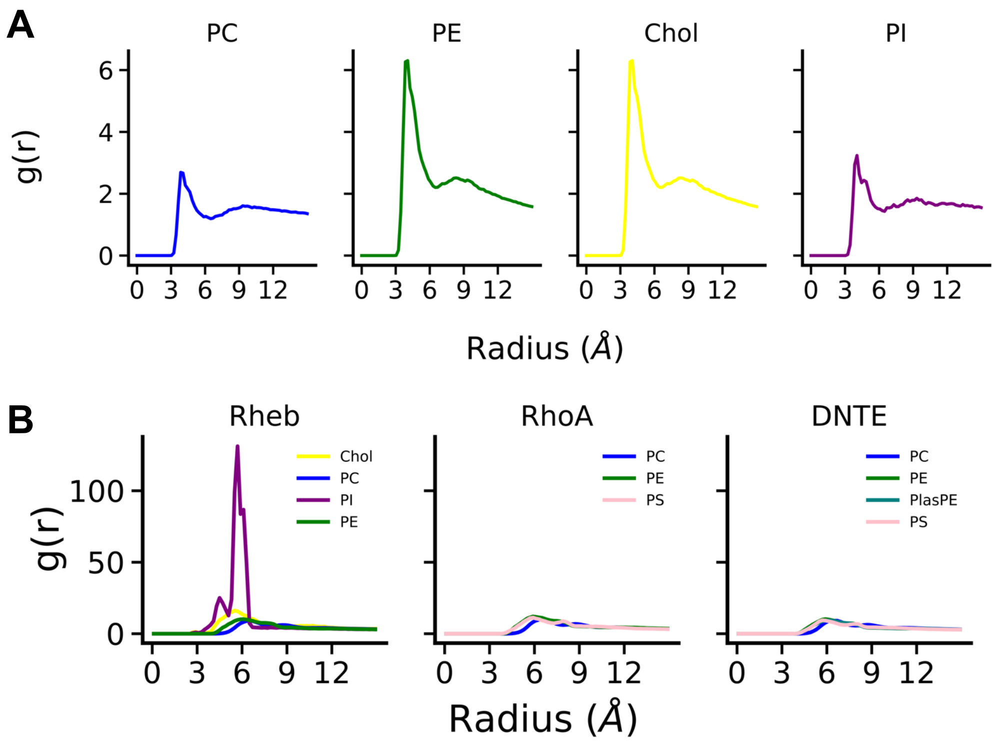


**Figure S7:** (**A**) Radial pair distribution function of all sidechain atoms on the Rheb peptide against lipid phosphorous atoms (PE, PI, PC) or hydroxyl oxygens (Chol). (**B**) Radial pair distribution function describing lipid-lipid interactions in the three systems, with pair distributions calculated using phosphate atoms (PE, PI, PS, PC, PlaPE) or hydroxyl oxygens (Chol)

### References


(1) Liang, X.; Jung, S. Y.; Fong, L. W.; Bildik, G.; Gray, J. P.; Mao, W.; Zhang, S.; Millward, S. W.; Gorfe, A. A.; Zhou, Y.; Lu, Z.; Bast, R. C. Membrane Anchoring of the DIRAS3 N-Terminal Extension Permits Tumor Suppressor Function. *iScience* **2023**, *26* (11), 108151. https://doi.org/10.1016/j.isci.2023.108151.

(2) Cui, H.; Lyman, E.; Voth, G. A. Mechanism of Membrane Curvature Sensing by Amphipathic Helix Containing Proteins. *Biophys. J.* **2011**, *100* (5), 1271–1279. https://doi.org/10.1016/j.bpj.2011.01.036.

(3) Drin, G.; Casella, J.-F.; Gautier, R.; Boehmer, T.; Schwartz, T. U.; Antonny, B. A General Amphipathic α-Helical Motif for Sensing Membrane Curvature. *Nat. Struct. Mol. Biol.* **2007**, *14* (2), 138–146. https://doi.org/10.1038/nsmb1194.

(4) Giménez-Andrés, M.; Čopič, A.; Antonny, B. The Many Faces of Amphipathic Helices. *Biomolecules* **2018**, *8* (3), 45. https://doi.org/10.3390/biom8030045.
